# Midline Thalamic Neurons Show Early Hypersynchrony During Absence Seizures in Mouse Models

**DOI:** 10.64898/2025.12.19.695246

**Authors:** Elzbieta Dulko, Scott Kilianski, Anna Grace Carns, Arya Rajesh, Isabelle Lesmana, Magdalena Pikus, Mark Beenhakker

## Abstract

Generalized seizures reflect a pathological state of sudden, global neuronal hypersynchrony. The mechanisms that support the initiation and maintenance of such synchrony remain unknown. Using simultaneous single-unit and electrocorticographic recordings in two mouse models of absence epilepsy, we evaluated the activity of approximately 2,000 individual neurons across 26 brain structures. By doing so, we resolved the temporal progression of single neuron activity prior to and during generalized seizures. Surprisingly, we observed that rhythmic, synchronized activity emerges early and gradually in both thalamus and cortex. Moreover, while we observe that individual neurons across most structures fire rhythmically and synchronously during seizures, a small subset of midline thalamic nuclei display activity that is strongly correlated across multiple regions. Finally, disturbing neuronal activity within the midline thalamus terminates seizures. Thus, our findings collectively highlight the possible role of the midline thalamus in emergence of neuronal hypersynchrony.

## Introduction

Neuronal synchrony is a key feature of the healthy brain and supports important functions such as sleep, attention, and motor coordination. Aberrant neuronal synchrony is often observed during pathological conditions, including schizophrenia^1^, Parkinson’s disease^2^ and epilepsy^3^. Generalized seizures associated with many forms of epilepsy are defined by the abrupt onset of synchronous neuronal activity across many brain structures. However, how such pathological hypersynchrony emerges—or even the extent to which such hypersynchrony exists—remains unknown. One form of epilepsy defined by recurrent generalized seizures is absence epilepsy, a common pediatric form of the disease. In humans, absence seizures are associated with the rapid emergence of 3-Hz spike-wave discharges (SWDs) across multiple cortical structures that coincide with impaired awareness^4^. Although some gene mutations are associated with absence epilepsy^5^, a clear understanding of disease etiology is lacking.

Several animal models have been developed to study absence epilepsy, including genetic and pharmacological models. Genetic rodent models include the GAERS and WAG/Rij rats, as well as the Stargazer, Tottering, Lethargic and C3H/HeJ mice; also, in the mid-1900s cat models of absence epilepsy were developed^6^. Pharmacological models, including γ-hydroxybutyrate (GHB)-and penicillin-induced models, have also been used to probe network mechanisms underlying SWD generation. Although not yet systematically compared, the likelihood that distinct ictogenic etiologies exist among these diverse models appears high. Nonetheless, these models collectively share the presence of SWDs and similar pharmacological sensitivities with human absence seizures, and several decades of work have established that generalized absence seizures are produced by recurrent cortico-thalamocortical networks. Experiments performed on acute rat and ferret brain slices demonstrate that thalamic nuclei—especially lateral structures such as the lateral geniculate and ventrobasal nuclei—contain rhythmogenic circuits capable of generating hypersynchronous neuronal oscillations resembling those observed during absence seizures^7–9^. *In vivo* experiments performed in rat models of absence epilepsy indicate that neuronal field activity recorded in the cortex consistently precedes thalamic activity during the early stages of the SWD^10,11^. Collectively, these studies laid the foundation for what is today known as the *Cortical Focus Theory* of absence seizure generation, wherein it is proposed that a bout of excessive cortical activity engages cortico-thalamo-cortical circuits that support and recruit additional structures to ultimately produce highly rhythmic and widespread neuronal oscillations in the form of SWDs^12^.

The *Cortical Focus Theory* of absence epilepsy proposes that seizure initiation involves focal abnormalities within the somatosensory cortex that subsequently engage the broader thalamocortical network. This theory was initially supported by local field potential (LFP) recordings demonstrating focal cortical abnormalities proceeding generalized SWDs in rodent models. Subsequent studies using *in vivo* intracellular recordings from layer 5/6 pyramidal neurons in the somatosensory cortex of anesthetized GAERS rats further supported the involvement of cortical neurons in SWD initiation^11^. Human magnetoencephalography studies also provide evidence for focal cortical involvement in patients with absence epilepsy^13^. Notably, however, human EEG-fMRI studies suggest that the intralaminar thalamic nuclei may be involved in SWD initiation and/or early propagation^14^. Thus, discrepant conclusions regarding SWD initiation may reflect differences among animal models and humans, differences in recording methodologies, and/or differences in anesthesia conditions. Nonetheless, perturbation studies have demonstrated that manipulating somatosensory cortical activity can influence SWD generation, further supporting an important role for cortical circuits^15,16^. Finally, recent multichannel extracellular recordings in awake, behaving animals provide new insights regarding neuronal spiking activity across cortico-thalamocortical networks during SWDs^17,18^. Interestingly, these studies have revealed that mean firing rates in both cortical and thalamic populations decrease during SWDs relative to baseline activity, suggesting that increased synchrony, rather than elevated firing rate, may be a defining feature of absence seizures. Resolving if and/or how a reduction in firing rates triggers SWDs remains a challenge for the field.

Here, we aim to resolve the spatiotemporal dynamics of neuronal synchronization that emerge during SWDs. We analyzed the activity of individual neurons before, during, and after SWDs in two models of absence epilepsy: C3H/HeJ^19–21^ and Stargazer mice^22–24^. As expected, SWDs were marked by widespread neuronal hypersynchrony across multiple brain structures in both models. However, localized clusters of hypersynchronous activity appeared in select thalamic nuclei up to 500 milliseconds before SWD onset (as defined by the motor cortex ECoG), suggesting that thalamic recruitment begins earlier than previously appreciated within the framework of the *Cortical Focus Theory.* Moreover, although SWDs recorded at the cortical surface in both models showed similar spectral features, the temporal evolution of the underlying neuronal dynamics differed, suggesting that circuit-level drivers of SWDs differ between models. Consistent with this idea, high-frequency stimulation of the midline thalamus disrupted ongoing SWDs, with differing effects in the two mouse models. Collectively, our results provide a novel perspective on SWD generation. *First*, changes in neuronal synchrony—not firing rates—are key hallmarks of SWDs. *Second*, the neural circuits recruited during SWDs are widespread, including those in several under-reported thalamic and hippocampal structures. *Finally*, SWD networks appear degenerate: although C3H/HeJ and Stargazer mice both produce electrographically similar SWDs, their single neuron spatiotemporal dynamics across structures differ.

## Results

### Experimental approach

Studies of absence seizure generation have investigated the contributions of numerous cortical and thalamic structures, including the somatosensory cortex (S1)^10,11^, reticular thalamus (RT)^25–27^, ventrobasal thalamus (VB)^17,18^, posterior nucleus (PO)^28,29^, anterior thalamic nuclei^30^, and intralaminar thalamic nuclei^31–34^. Despite these findings, the prevailing thalamocortical network model continues to emphasize interactions among S1, RT, and VB as the core circuitry underlying SWD generation^17,18^. We evaluate this model in the C3H/HeJ and Stargazer mouse. We also examine additional thalamic nuclei located near the brain’s midline and the hippocampus, a structure with only a limited understanding of SWD engagement^35–38^.

We simultaneously recorded electrocorticographic (ECoG) signals and single-unit activity in awake, head-fixed C3H/HeJ and Stargazer mice. Representative recordings demonstrate that SWDs were observed across most brain structures, albeit with variable prominence (**Fig. 1a**). A representative raster plot of single-unit activity during one C3H/HeJ SWD is shown in **Fig. 1b**. Silicon probe location was verified with immunofluorescence and systematic reconstruction of the resultant fluorescent DiI tracks (**Fig. 1c**). Our dataset (**Fig. 1d**) includes single-unit activity from 433 SWDs across 26 structures in 28 C3H/HeJ mice (14 M, 14 F) and 1521 SWDs across 13 brain structures in 21 Stargazer mice (6 M, 15 F). Each recording session yielded simultaneous measurements from tens of neurons (median = 25 and 33 for CH3/HeJ and Stargazer mice, respectively). Complete structure names and sample sizes are provided in **Table S1**. Despite similar electrographic appearances (**Fig. 1e**), Stargazer mice had 1.21 ± 0.74 SWDs/min whereas C3H/HeJ mice had 0.38 ± 0.39 SWDs/min (p<0.0001, unpaired t-test). Additionally, Stargazer SWD duration was prolonged (5.15 ± 1.5 sec), relative to C3H/HeJ SWD duration (3.88 ± 1.35 seconds, p<0.01, unpaired t-test). Prolonged SWD duration reflected a larger number of spike-and-wave cycles per Stargazer SWD (29.30 ± 8.36 cycles), relative to C3H/HeJ SWD (21.20 ± 6.04 cycles, p<0.001, unpaired t-test). An SWD cycle was defined as the interval from one prominent, negative trough to the next. The average SWD cycle duration was equivalent between Stargazer (0.18 ± 0.02 sec) and C3H/HeJ mice (0.18 ± 0.02 sec, p=0.24, unpaired t-test, **Fig. 1f**).

**Figure 1.**
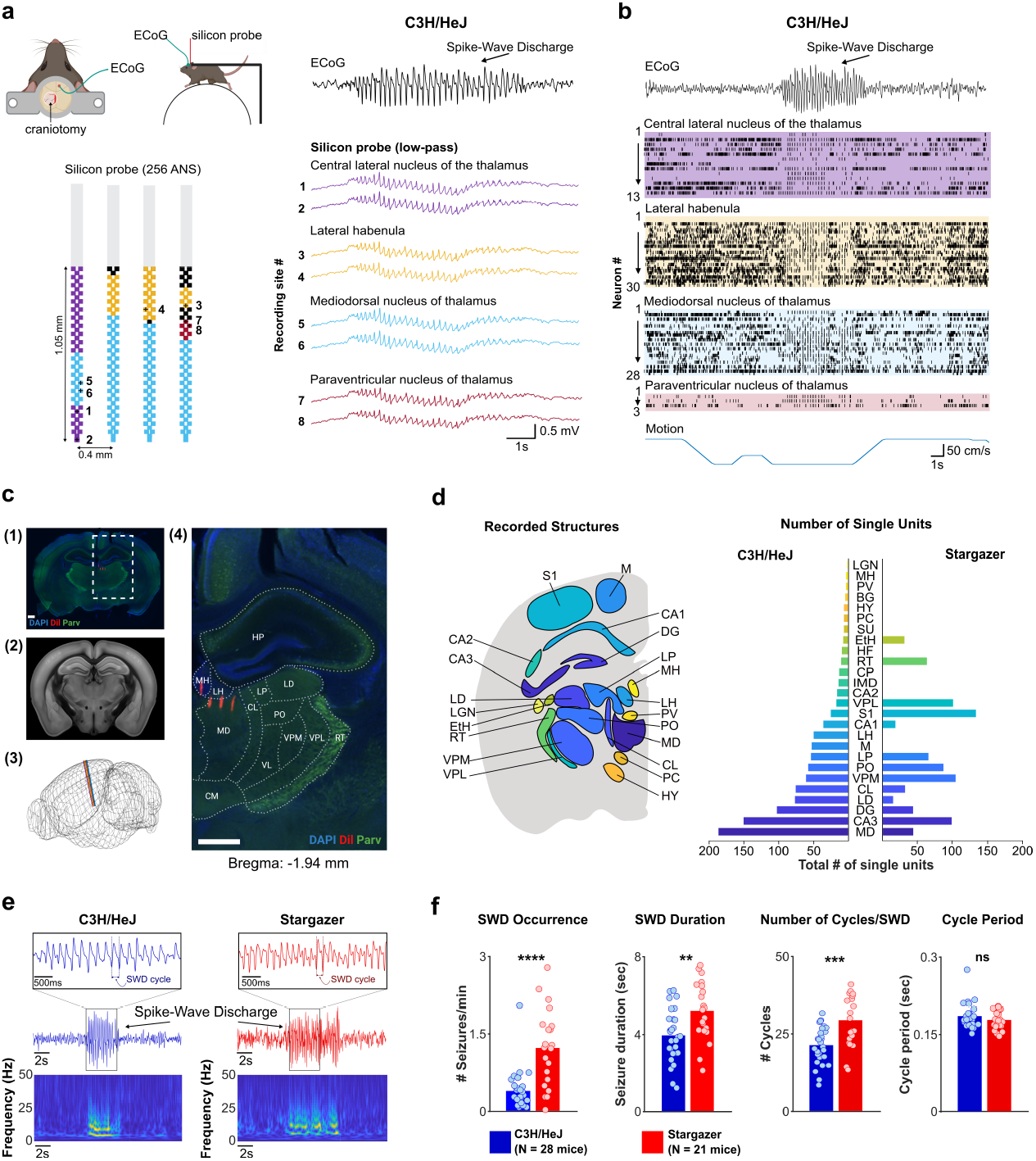
Approach and summary of collected data. *a*. Mice were implanted with headplates and ECoG electrodes (top). Seven days after headplate implantation, mice were head-fixed to the recording apparatus and recorded. Schematic of the multi-electrode silicon probe (left) and a representative ECoG and low-pass silicon probe recording of one SWD (right). Recordings are color-coded according to the electrode position on the silicon probe. SWDs were detected across most recorded brain structures, although their amplitude varied between regions ***b.*** The action potential activity of 74 neurons is shown in raster format. Bottom trace shows mouse motion. Note behavioral arrest during the SWD. ***c*.** After the recording, brains were fixed and sectioned. Sections with visible probe tracks (1, 4) were aligned with the Allen Institute Brain CCF Framework ^78^(2) to verify probe location (4). Scale bar, 500 µm. ***d*.** Recorded brain structures (left) and total number of single units recorded in each brain structure for C3H/HeJ and Stargazer mice (right). BG, SU, HF, CP are not shown in left panel. ***e.*** Representative SWD and spectrogram from a C3H/HeJ and Stargazer mouse. Insets show an expanded view of the ECoG trace, illustrating the characteristic spike-and-wave morphology. A single SWD cycle, defined from negative trough to the next, is marked. ***f.*** Comparison of seizure features between C3H/HeJ and Stargazer mice (bars represent means; unpaired two-tailed student’s t-test, **** p<0.0001, *** p<0.001, ** p<0.01, n = 28 C3H/HeJ mice, n = 21 Stargazer mice). Schematic in (***a***) and (***c***) created with BioRender.com. See Table S1 for complete structure names and sample sizes.

### Neuronal firing during SWDs

We first quantified single unit firing rates during SWDs (**Fig. 2**). This analysis revealed different response profiles across nuclei. Neuronal firing rates in some nuclei decreased during SWDs (e.g., VPM, **Fig. 2a**, left), whereas neuronal firing rates in other nuclei increased (e.g., lateral posterior nucleus of thalamus, LP, **Fig. 2a**, right). We compared mean firing rates during SWDs to those during non-seizure baseline periods, defined as a duration-matched, non-seizure epoch positioned midway between two consecutive SWDs (**Fig. 2b**). We observed that neuronal firing in the somatosensory cortex and VPM generally dropped during the SWD relative to non-seizure periods, observations consistent with previous studies that evaluated firing rates within the canonical thalamocortical network^17,18^. A similar trend was observed in RT (**Fig. 2c, Table S2**). We also evaluated single unit firing rates among structures beyond the canonical thalamocortical structures implicated in SWD generation. In the vast majority of structures, we observed only modestly lower firing rates during the SWD, relative to non-seizure periods (**Fig. 2c, Table S2**).

**Figure 2.**
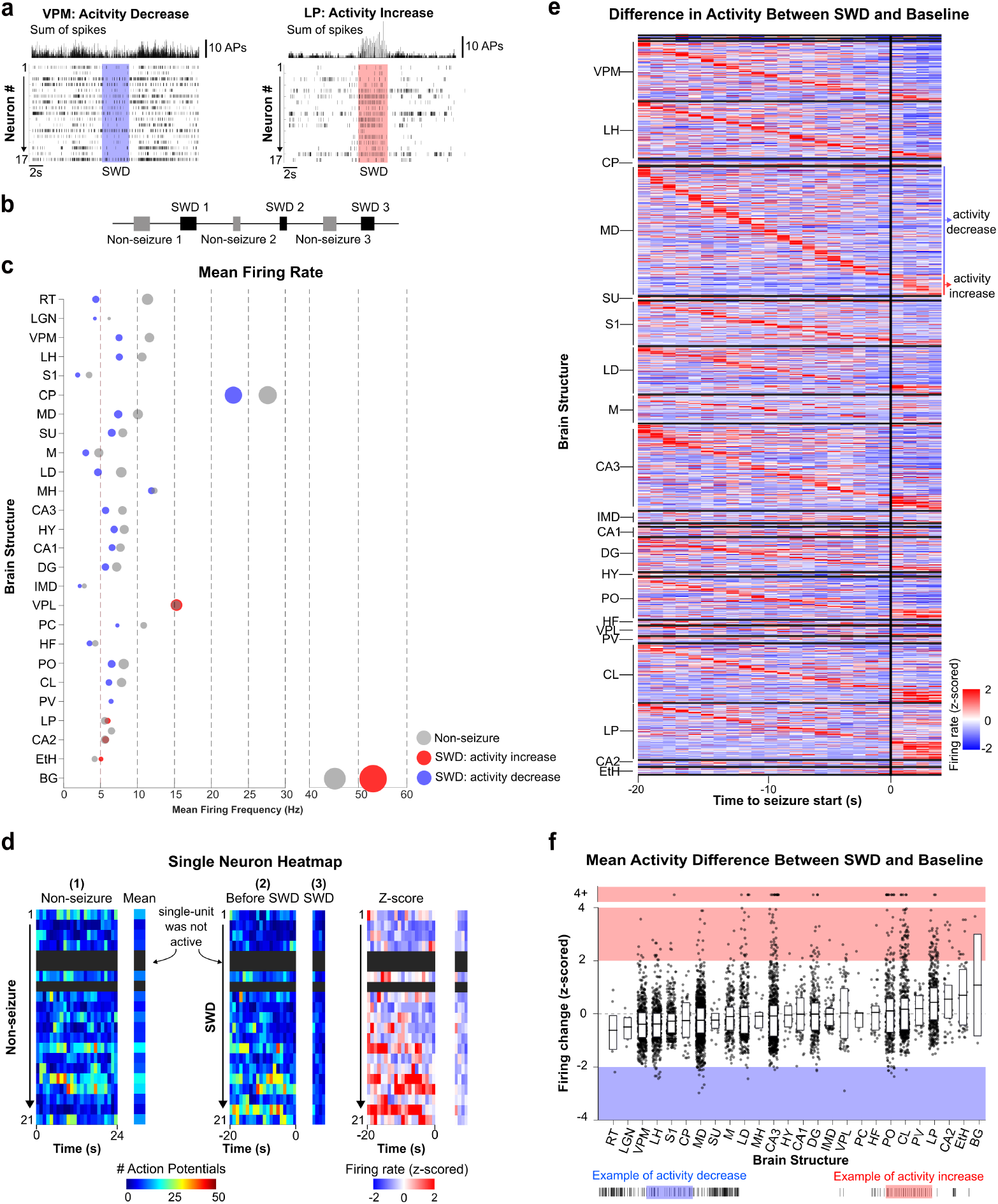
SWDs produce only subtle changes in mean firing rate. ***a***. Action potential activity of individual neurons shown in raster format (bottom) and overall spike count (top). Shown are neuronal firing activities in ventral posteromedial thalamus (left) and lateral posterior thalamus (right) during one SWD. The shaded area indicates SWD occurrence. ***b***. Firing rates of neurons during SWDs (black) were compared to firing rates during non-seizure periods (gray). ***c***. Dot plot showing mean firing rate for non-seizures and seizures across all brain structures. Dot size is scaled based on standard deviation. Blue and red dots correspond to structures wherein activity levels trended lower or higher, respectively, during the SWD versus non-seizure periods. ***d***. Z-scored heatmap for a single neuron recorded in one mouse. Z-scores were calculated for each 1-second time bin before (2) and during SWD (3), relative to the mean baseline firing computed during the corresponding non-seizure event (1). ***e***. Z-scored heatmap showing firing rate differences between seizure and non-seizure periods for neurons of each brain structure. Each row represents the z-scored firing activity of a single neuron during an individual SWD (neuron-SWD pair). Rows are grouped by brain structure and sorted according to the timing of the peak z-score, with neurons exhibiting earlier peak activity positioned toward the top and neurons exhibiting later peak activity positioned toward the bottom. This visualization highlights the heterogenous timing and magnitude of neuronal responses associated with SWDs. RT, LGN, HF, MH, BG and PC are not included in the heatmap because these structures contained too few single units. ***f***. Box plot of mean firing change for each brain structure. Each dot represents one neuron during one seizure. The bottom and top of each box correspond to the 1^st^ (Q1, 25^th^ percentile) and 3^rd^ quartile (Q3, 75^th^ percentile), respectively; the line inside the box indicates the mean. Blue and red rectangles denote z-score <-2 and=2 respectively. n = 28 C3H/HeJ mice. See Table S1 for complete structure names and sample sizes.

To better resolve the temporal progression of activity leading up to a seizure, we examined the z-scored firing rates of individual neurons leading to—and during the first four seconds of—many consecutive SWDs. The example neuron shown in **Fig. 2d** generally decreased its activity during each of the 21 seizures. We then visualized z-scored firing rates by grouping them by brain structure (**Fig. 2e**). Each row represents one neuron during one SWD, with rows ordered according to peak firing activity. This analysis revealed a spectrum of SWD-associated changes in neuronal activity levels in most structures. Most neurons exhibited insignificant changes in firing during SWDs; however, elevated firing rates were observed in a subset of neurons (e.g., mediodorsal thalamus, MD, **Fig. 2e**). Nonetheless, in aggregate, the mean neuronal firing rates (z-scored) during the SWD were not significantly different from non-seizure activity levels, across all structures (**Fig. 2f**). To confirm that the observed firing rate trends did not result from a biased approach to only evaluate algorithmically sorted units, we also quantified firing changes using the entire spiking activity (ESA) method^39^ (see *Methods*). ESA analysis confirmed a general decrease in mean firing activity during SWDs (**Fig. S7e**).

To assess whether the firing patterns we observed in C3H/HeJ mice are mouse model-specific, we quantified firing rates of single units in the Stargazer mouse, a long-standing model of absence epilepsy^22–24^. Similar to C3H/HeJ mice, we found that Stargazer neurons also exhibited variable firing rate changes associated with the SWD (**Fig. S1a-d**). Whereas the mean neuronal firing rate within most Stargazer brain structures was modestly lower during SWDs compared to non-seizure periods, neurons within a small subset of Stargazer structures appeared to fire more strongly during SWDs. Although changes in firing rates across Stargazer structures did not reach statistical significance (i.e., mean z-scores between-1 and 1), we observed a few opposing trends in some brain structures, suggesting model-specific differences. For example, firing in the LP nucleus increased during SWDs in C3H/HeJ mice (mean z-score > 0; **Fig. 2f**) but decreased in Stargazer mice (mean z-score < 0; **Fig. S1d**).

Although the overall mean population firing rate during SWDs did not differ from baseline, a subset of units in some structures increased their firing rates during SWDs (**Fig. 2e, f**). Thus, the absence of a population-wide change in firing does not preclude changes in activity among specific neuronal subpopulations, consistent with previous reports that seizure-related firing changes are often restricted to a minority of recorded neurons rather than observed uniformly across structures^18,40^. To further characterize the identity of units exhibiting SWD-related firing changes, we classified units into putative excitatory and putative inhibitory populations based on extracellular waveform features^41^ (*see* Methods “Unit classification”, for details; **Fig. S1e, f**). For each brain structure and cell type, we identified units exhibiting consistently elevated firing rates during SWDs, defined as a mean z-scored firing rate > 2 for at least 50% of SWDs (**Table S3**). In C3H/HeJ mice, consistently elevated firing was observed in both putative excitatory and inhibitory populations, although the largest fractions were generally observed among putative excitatory units, including CL (13/63 units, 21%), LP (11/51 units 22%, CA3 (15/97 units, 15%). Across structures, consistently responsive excitatory units typically represented a minority of recorded units (median, 4%, range: 0-43%). In Stargazer mice, fewer units met our criteria for consistently elevated firing rates during SWDs, a likely reflection, at least in part, of the larger number of analyzed SWDs per neuron and/or greater seizure-to-seizure variability in the mice. Overall, these findings indicate that SWD-related elevated firing rates were generally restricted to a subset of recorded units.

Thalamic neurons are commonly defined by their ability to fire repetitively at a modest rate (i.e., “tonic mode”) or in brief bursts of action potentials generated at a high rate (i.e., “burst mode”)^26,42,43^. Classically, SWDs are associated with thalamic neurons firing in burst mode^7,44^. However, across most structures in both C3H/HeJ and Stargazer mice we found that neuronal firing was not obviously biased towards burst mode during the SWD, a conclusion that is supported by the apparent robustness of spike sorting algorithms to detect rapidly firing single units despite subtle changes in spike morphology (**Fig. S2c**). To this end, we first calculated the instantaneous firing rate of each C3H/HeJ neuron during SWDs and non-seizure periods. We then defined neuronal activity as tonic or bursting based on whether the instantaneous firing rate was less than or greater than 200 Hz, respectively. Only in a few structures did we observe a subtle increase in the proportion of bursting neurons during the SWD, relative to non-seizure periods, indicating that burst mode is not overwhelmingly favored during the SWD (**Fig. S2b**). Even in brain structures wherein burst firing was slightly elevated during SWDs (e.g., hippocampal CA1, CA1; ventral posterolateral thalamus, VPL; VPM, posterior complex of thalamus, PO), most neurons did not burst on each cycle of the SWD (**Fig. S2d, e**), an observation consistent with recent work^17^. We also quantified burst firing in Stargazer mice and similarly found only a modest increase in most brain structures (**Fig. S2f**). Together, these findings indicate that SWDs are not associated with a widespread increase in neuronal firing rate but rather likely involve heterogeneous firing responses in which specific neuronal subpopulations exhibit elevated activity patterns.

### Oscillatory activity of neuronal population firing during SWDs

Although the firing rates of many individual neurons changed during SWDs, such changes were generally subtle and highly variable within a structure (**Fig. 2e, f**). Thus, when viewed through the lens of neuronal firing rates, it becomes difficult to resolve if any one structure initiates and/or maintains a SWD. We therefore evaluated the oscillatory nature of neuronal firing, as the most salient feature of the single units we recorded—across nearly every structure—was the rapid entrainment to the large, negative-going spike of the SWD. Since SWDs are generalized events, it is presumed that participating brain structures generate rhythmic firing patterns that follow the dominant rhythm of the cortical ECoG during the SWD (**Fig. 3a, b**). To test this possibility, we examined the coupling between neuronal firing and the cortical ECoG using spike-field coherence (SFC) analyses^45^. SFC is a frequency domain measure of the linear association between two continuous time series^46^; in our case, these time series correspond to the ECoG and the time-varying spike count. We restricted our SFC analysis to the duration of the SWD, which revealed that neuronal populations in most recorded brain structures produce oscillatory activity with a peak in the 5-6 Hz range, a frequency that aligns with the dominant frequency associated with C3H/HeJ and Stargazer SWDs. Indeed, the peak of coherence in the 5-6 Hz range suggests a tight relationship between the ECoG signal and neuronal firing (C3H/HeJ: **Fig. 3c, d**; Stargazer: **Fig. S3a**). We next represented mean coherence at all frequencies for all recordings as a heatmap (**Fig. 3f**, left). To examine whether neuronal activity in each brain structure aligns with the dominant seizure frequency, we plotted a heatmap of only maximum coherence frequencies, revealing that most brain structures are coupled to the 5-6 Hz rhythm (**Fig. 3f**, right). In a few brain structures (e.g., centrolateral thalamus, CL; MD, LD), neurons tended to fire in phase with the SWD cycle, but they also fired outside of the dominant oscillatory cycle, resulting in weaker phase-locking and resulting in peak coherence in 1-2Hz range (**Fig. 3f**, right). Additionally, we evaluated neuronal rhythmicity using the ESA approach (**Fig. S5**). Consistent with the aforementioned SFC analysis, ESA revealed that PO, VPL, CL, and MD, all thalamic nuclei, exhibited the highest increase in neuronal rhythmicity during the SWD, relative to non-seizure periods.

**Figure 3.**
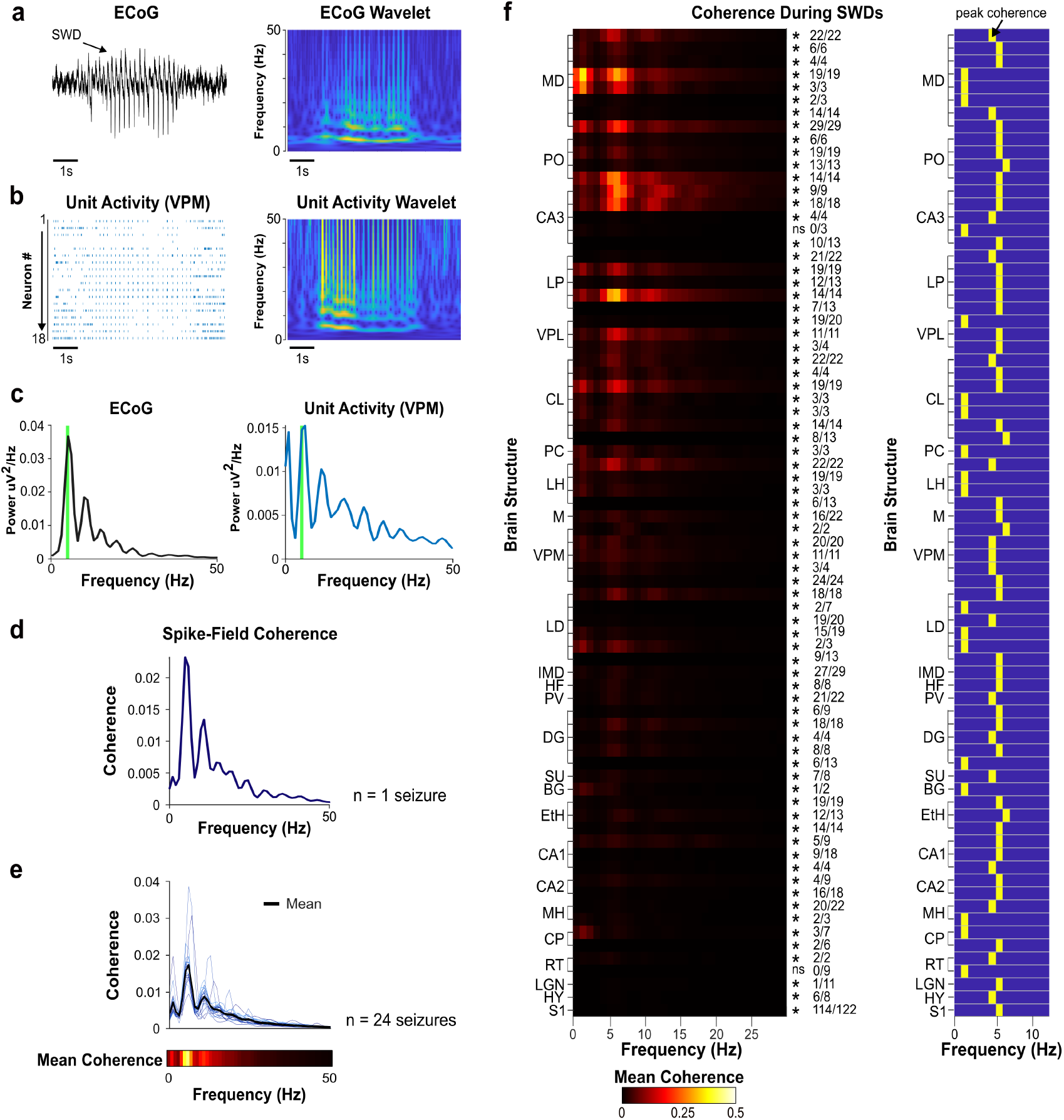
5 Hz oscillatory activity exists in most recorded C3H/HeJ brain structures. ***a***. ECoG (left) and corresponding frequency wavelet spectrogram (right). ***b***. Raster plot of firing activity in the thalamic VPM nucleus (left) and frequency wavelet spectrogram based on the population firing count (right). ***c***. Power spectrum for ECoG (left) and VPM population firing count (right). For reference, the green line is placed at 5.8 Hz. ***d***. Calculated spike-field coherence (SFC) for one SWD. ***e***. SFC for all individual SWDs (thin lines) and population mean (thick line). The linear heatmap below represents the mean SFC across all SWDs, n = 1 C3H/HeJ mouse. ***f.*** Heatmap of mean SFC values across all SWDs in one recording organized by the brain structure (left). Asterisks (*) mark rows where SFC in the 4.5-6.5 Hz band was statistically significant. Fractions (e.g., 22/22) show the number of statistically significant SWDs over the total number of SWDs (Surrogate-based significance test, *, p<0.05. n = 28 C3H/HeJ mice). A complementary heatmap of peak SFC values highlights a consistent maximum at 5 Hz (right).

### Spatiotemporal map of firing before, during, and after SWDs

Spike-field coherence analysis measures the coupling strength between neuronal firing and ECoG signals at specific frequencies. We next sought to determine the precise timing of action potentials within individual SWD cycles. Therefore, we defined action potential times in terms of a normalized cycle period, an approach commonly used to reveal neuronal activity patterns associated with rhythmically active neural circuits^47^. A cycle was defined as the period between one negative-going SWD spike in the ECoG (phase = 0) and the next negative-going spike (phase = 1; e.g., **Fig. 4a**). Normalizing action potential times in this manner enables the aggregation of neuronal activity over multiple seizures and across animals despite variability in the spectral composition of the SWDs. Our phase analysis of C3H/HeJ SWDs revealed a clear temporal sequence of action potential firing during the SWD (**Fig. 4c, e**). Notably, leading up to the negative, ECoG spike during an ongoing SWD, synchronous neuronal activity is evident in several structures, including VPM, LD, and hippocampal CA3 (CA3). Also notably, these structures are active before S1 (**Fig. 4c** and **Fig. S8**). Neurons of the basal ganglia (BG) and caudate putamen (CP) were not phase-locked during SWDs observed in C3H/HeJ mice (**Fig. 4c**, BG: p=0.051, CP: p=0.66). This analysis also revealed that activity within the RT nucleus occurs mostly after the negative SWD spike, during the wave component, a finding that is consistent with previous studies^48^. To further examine whether neuronal identity influenced dynamics during SWDs, we incorporated cell-type classification into our phase-locking analysis (**Fig. 4c**, right; **Fig. S3b**). Across brain structures, both excitatory and inhibitory populations generally fired around the negative ECoG spike, with only modest differences in preferred firing phase. In most structures, putative excitatory units tended to precede putative inhibitory units by a small fraction of the SWD cycle.

**Figure 4.**
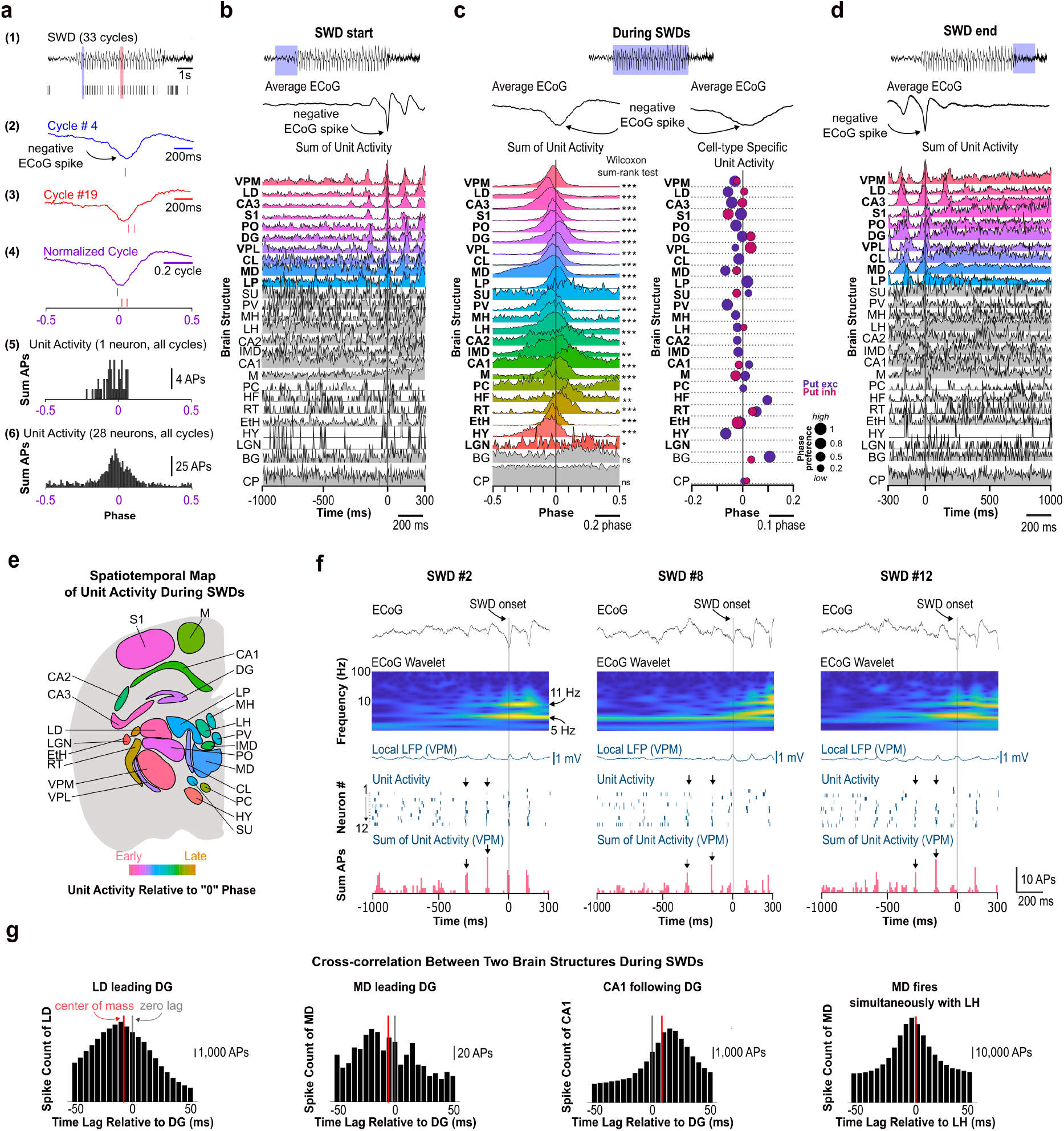
Correlated firing activity before, during, and after SWDs. ***a***. Phase analysis. (1) ECoG signal during a seizure and action potential raster from a single neuron in MD. Two cycles are highlighted in blue (cycle 4) and red (cycle 19). (2) Expanded view of the blue cycle and neuronal firing relative to this cycle. (3) Expanded view of the red cycle and neuronal firing relative to this cycle. (4) Neuronal firing during the blue and red cycles relative to a normalized cycle. (5) Histogram of firing during all 33 cycles, n = 1 neuron. (6) Histogram of firing during all 33 cycles for all neurons, n = 28 neurons. ***b***-***d.*** Histograms of neuronal firing for all brain structures before (***b***), during (***c***), and following (***d***) the SWD. Data from all mice are represented in a single histogram. Scatter plot in panel **c** shows phase preference for putative excitatory and inhibitory neurons; dot size indicates the magnitude of phase-locking. Brain structures in panels ***b***-***d*** are arranged by the onset of rhythmic spiking observed before SWD (panel ***b***). Spike count was normalized to the peak firing (Wilcoxon rank-sum test, *** p<0.001, ** p<0.01, * p<0.05. n = 28 C3H/HeJ mice). ***e***. Graphical representation of the order of firing during seizures. Color denotes when the peak firing occurs (based on panel ***c***). ***f***. VPM neurons oscillated before SWD onset. Synchronous firing, shown in the raster and spike counts (black arrows), emerged hundreds of milliseconds before SWD onset. Wavelet analysis of the ECoG confirmed no increase in 5 Hz power prior to SWD onset, ensuring accurate SWD detection. Despite high unit synchrony, the local LFP measured in VPM did not always reveal prominent peaks. ***g.*** Representative histograms of the cross-correlation between all pairs of neurons from distinct brain structure pairs during SWDs. Cross-correlations are shown over a ± 50 ms window with 5 ms bin size. The gray vertical line marks zero lag, and the red line indicates the center of the mass (average lag) of each histogram. Order of firing is consistent with phase analysis in panel ***c***. For all cross-correlations, see Fig. S6.

To validate whether phase relationships observed in single-unit activity were also reflected in unsorted data population activity, we performed an analogous phase-locking analysis on the ESA recorded from each structure in C3H/HeJ mice. Consistent with our single-unit data, ESA revealed weak phase-locking in BG and CP, and strong phase-locking in PO, EtH, and VPL (**Fig. S8d, e**). Furthermore, the temporal order revealed by cross-correlation measures among pairs of neurons recorded simultaneously during a single experiment (**Fig. 4g** and **Fig. S6)** closely matched that obtained from our phase analyses (c.f., **Fig. 4c**), indicating that our observed temporal sequence across structures is consistent across analytical approaches and independent of the chosen reference signal.

Notably, we found that about a third of all brain structures recorded in the C3H/HeJ mouse contain rhythmic spiking (i.e., repeating peaks spaced ∼180ms apart in **Fig. 4b** and **4d**) prior to the start of the SWD. This finding suggests that although SWDs are rapidly generalized events, the events begin only within a subset of brain structures (i.e., top, colored structures in **Fig. 4b**). Brain structures that produced rhythmic activity *before* the SWD also produced rhythmic activity *after* the SWD. Also, we found that rhythmic single unit activity within SWD-leading structures was not associated with salient signals in the local LFP (i.e., the LFP measured by the electrode placed within the recorded structure), or the cortical ECoG (**Fig. 4f** and **Fig. S3a, b**). This observation suggests that the LFP is not always a reliable indicator of locally organized spiking activity in SWD-leading structures. This observation also suggests that local, organized spiking activity in these structures occurs without overt synaptic drive.

To determine if the spatiotemporal progression of neuronal activity we observed in the C3H/HeJ mouse is unique among mouse models, we evaluated phase relationships of single units (**Fig. S3b**) and ESA (**Fig. S7b**) in the Stargazer mouse. This analysis revealed several similarities between C3H/HeJ and Stargazer mice. First, we found that most brain structures were phase-locked to the ECoG signal during C3H/HeJ and Stargazer SWDs (**Fig. S3b**). However, peak neuronal firing occurred at slightly different periods within the C3H/HeJ SWD cycle relative to the Stargazer SWD cycle (**Fig. S3c**, e.g., S1, CL); nonetheless, offsets of no more than 0.2 phase units (∼40 ms) were observed. The presence of rhythmic activity in the Stargazer mouse (**Fig. S7b**) suggests that synchrony among thalamic structures (e.g., CL, MD, and PO) occurs gradually prior to cortical SWD onset in both mouse models.

### Neuronal synchrony during SWDs

SWDs manifest as high amplitude events in the ECoG signal. High amplitude events observed in LFP recordings (e.g., ECoG) generally occur when synaptic activity produced by a sufficiently large neuronal population occurs synchronously^49^. As such, the synchronization of neuronal action potentials that ultimately generate synaptic activity can only be indirectly inferred from LFP recordings. Notably, the *Cortical Focus Theory* was derived largely from LFP recordings^50^. Thus, after establishing that neurons fire rhythmically in many structures (**Fig. 3**) and are phase-locked to the ECoG SWD (**Fig. 4**), we next aimed to directly resolve the extent of neuronal synchronization among active neurons within the same structure, as well as across different structures.

We evaluated neuronal synchrony by calculating the Pearson correlation coefficient derived from spike trains obtained from pairs of recorded neurons^51,52^ (**Fig. 5a**). We first evaluated pair-wise, correlated activity between neurons that reside within the same brain structure (i.e., intra-nucleus correlations) using a bin size of 25 ms (**Fig. 5b1**); high correlation values result when both neurons fire coincidently within 25 ms. Importantly, pair-wise correlations were only weakly impacted by firing rates (r^2^ = 0.07; **Fig. 5b2**). We then measured changes in pair-wise neuronal synchrony during SWDs relative to non-seizure periods (i.e., correlation during SWDs – correlation during non-seizure periods). Not surprisingly, we found that the degree of synchrony increased during SWDs for most neuronal pairs recorded within the same structure (**Fig. 5a**). However, the extent of elevated, pair-wise synchrony varied across different nuclei (**Fig. 5b**). Notably, the largest increases in intra-nucleus synchrony were observed in thalamic nuclei in both C3H/HeJ (**Fig. 5b1**) and Stargazer mice (**Fig. S3e**).

**Figure 5.**
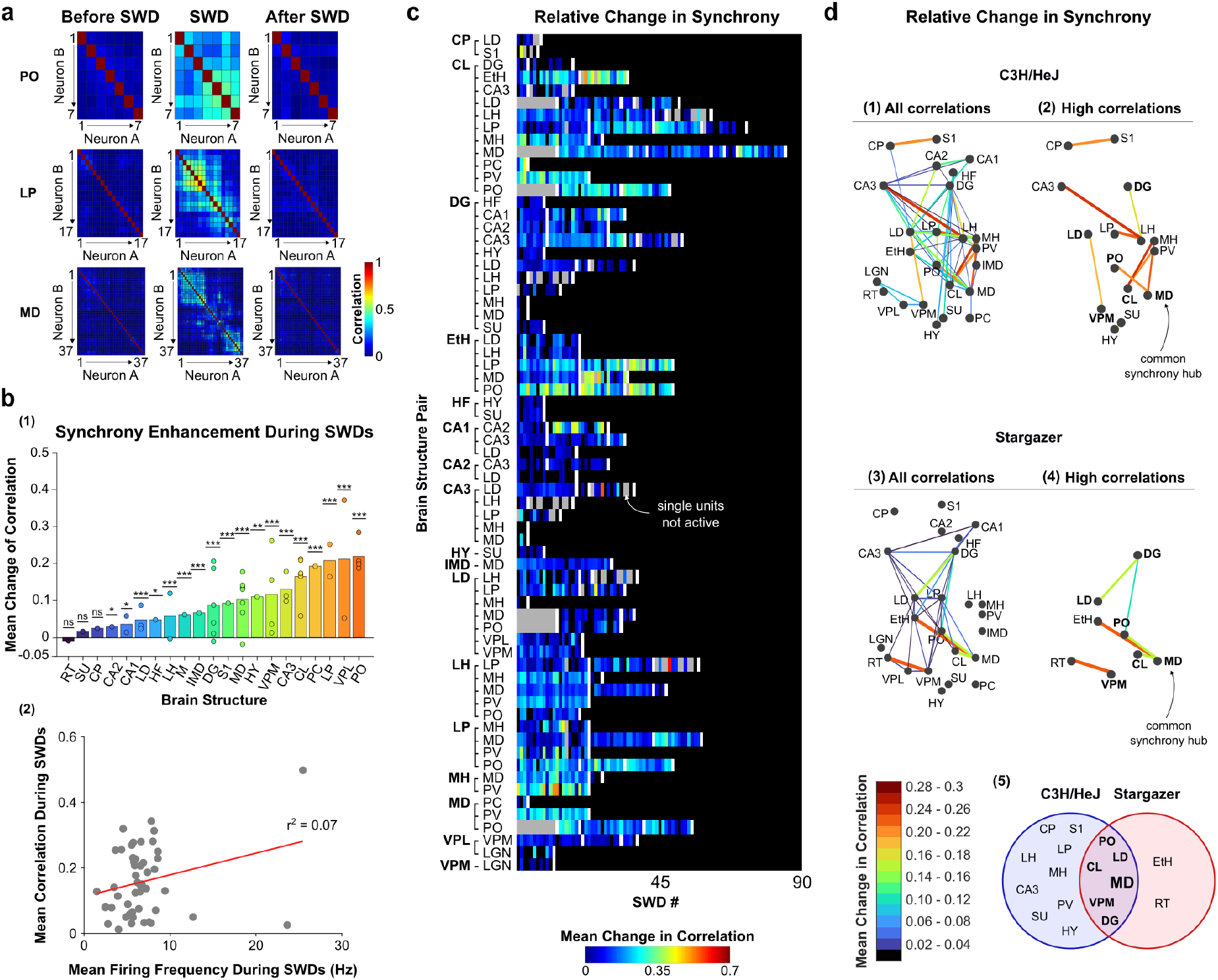
Increased neuronal synchrony during SWDs. ***a***. Pearson correlations reveal increased neuronal synchrony within a structure during SWDs. ***b***. (1) Increased neuronal synchrony *within* structures during SWDs, relative to baseline. Bars are sorted according to the mean increase in correlation values. Dots represent different mice. Statistical comparisons were made using linear mixed models to account for mouse-specific effects. Asterisks represent statistical significance in pairwise post hoc two-sided comparisons: *** p<0.001, ** p<0.01, * p<0.05. (2) A weak correlation exists between mean firing rates and the mean Pearson correlations during SWDs. Dots represent different brain structures. ***c***. Pearson correlations reveal increased neuronal synchrony *between* two structures during SWDs. The heatmap shows the mean increase in correlated spiking activity between two neurons recorded in two different structures during the SWD, relative to baseline (time-matched non-seizure event). Each distinct bar in the heatmap represents an average correlation increase for all pair-wise comparisons during one SWD. White bars demarcate SWDs from different mice. For example, the third row from the top of the heat map reveals correlated activity between neurons of the CL nucleus and the dentate gyrus. Such correlations were derived from two mice producing 3 and 13 SWDs, respectively. Gray bars indicate that single units were not active during the SWD and, therefore, correlation values were not calculated. ***d***. Functional connectivity measures derived from Pearson correlations, across brain areas recorded in the same recording session. Circles represent brain areas positioned by approximate anatomical location. Line width and color is scaled according to the correlation value. Panels 1,3: all correlations (no threshold); panels 2,4: high correlations (> threshold). See Fig. S4 for functional connectivity graphs across different correlation thresholds. Panel 5 shows a venn diagram of brain structures exhibiting high correlations in both models. n = 28 C3H/HeJ, n = 21 Stargazer mice. The bin size for panels ***a*** and ***b*** equals 25 ms, whereas for ***c*** and ***d*** bin size equals 50 ms. Panel ***a*** created with BioRender.com.

Next, we evaluated pair-wise synchrony between neurons located across different structures (i.e., inter-nucleus correlations) of C3H/HeJ mice; here, a bin size of 50 ms was used. For each SWD, pair-wise correlations were calculated over the duration of the seizure and then averaged across seizures. Changes in Pearson correlation values among pairs of recorded neurons during SWDs relative to non-seizure epochs are presented as a heatmap (**Fig. 5c**). Using graph theory^53^, these SWD-associated changes in correlated activity are presented as putative functional connections among structures (**Fig. 5d**). The effect of varying the correlation threshold on the resulting functional connectivity graphs is shown in **Fig. S4**. Pair-wise correlated activity increased during SWDs for all brain structure pairs (**Fig. 5d1**, all C3H/HeJ correlations) but increased most profoundly between CA3 and lateral habenula (LH) (mean = 0.28), as well as between CL and medial habenula, MH (mean = 0.27). Indeed, thalamic structures — particularly those along the midline thalamus — collectively exhibited the greatest increase in correlated activity (**Fig. 5d2**, high C3H/HeJ correlations). Thalamic structures recorded in the Stargazer mouse also exhibited the greatest increase in correlated activity during SWDs (**Fig. 5d3,4**, **Fig. S3f**). Altogether, lateral (LD, VPM) and midline (PO, CL, MD) thalamic brain structures, as well as the dentate gyrus (DG), were classified as having high correlations in both mouse models (**Fig. 5d5**).

Finally, to further identify brain structures potentially important for widespread synchrony (i.e., structures with high degree centrality), we performed degree centrality analysis, an approach used to resolve brain structures highly engaged within a broader neural network^53^; this analysis essentially identifies structures with a high number of functional connections. We identified MD thalamus as a common synchrony hub during SWDs, as this nucleus uniquely exhibited highly correlated activity with at least two brain structures in both mouse models. (**Fig. 5d2,4,5**). Thus, thalamic structures in both the C3H/HeJ and Stargazer mouse exhibit early signs of coordinated neural activity (**Fig. 4b,f; Fig. S3b**), produce the highest levels of neuronal synchrony (**Fig. 5a,b; Fig. S3e**), and appear to coordinate neural activity across multiple structures (**Fig. 5d**). Notably, however, degree centrality identifies structures with increased functional associations during SWDs but does not imply that these relationships are constant throughout seizure evolution or that they reflect directional interactions. Thus, future studies using time-resolved and directed connectivity approaches will be important to further define the dynamic organization of seizure networks.

### Frequency-dependent modulation of SWDs by the midline thalamus

We found that midline thalamic nuclei, including PO and CL, exhibited high levels of neuronal synchrony preceding and during SWDs (**Fig. 4b, c**). Additionally, we identified MD as a structure with high functional connectivity during SWDs in both C3H/HeJ and Stargazer mice (**Fig. 5d**). Indeed, as early as 1949, Hunter and Jasper proposed a role for the midline thalamic nuclei in SWD generation^6^, yet this prescient hypothesis has remained largely unexplored. Therefore, we next tested whether direct perturbation of midline thalamic neuronal activity would impact SWD generation in the C3H/HeJ and Stargazer models. We positioned stimulating electrodes such that current passed through the midline thalamus, allowing direct modulation of thalamic activity (**Fig. 6a1**). Correct electrode placement was verified by post hoc immunofluorescence (**Fig. 6a2**). Anatomical coordinates of electrode placements for all experimental mice are provided in **Table S4**.

**Figure 6.**
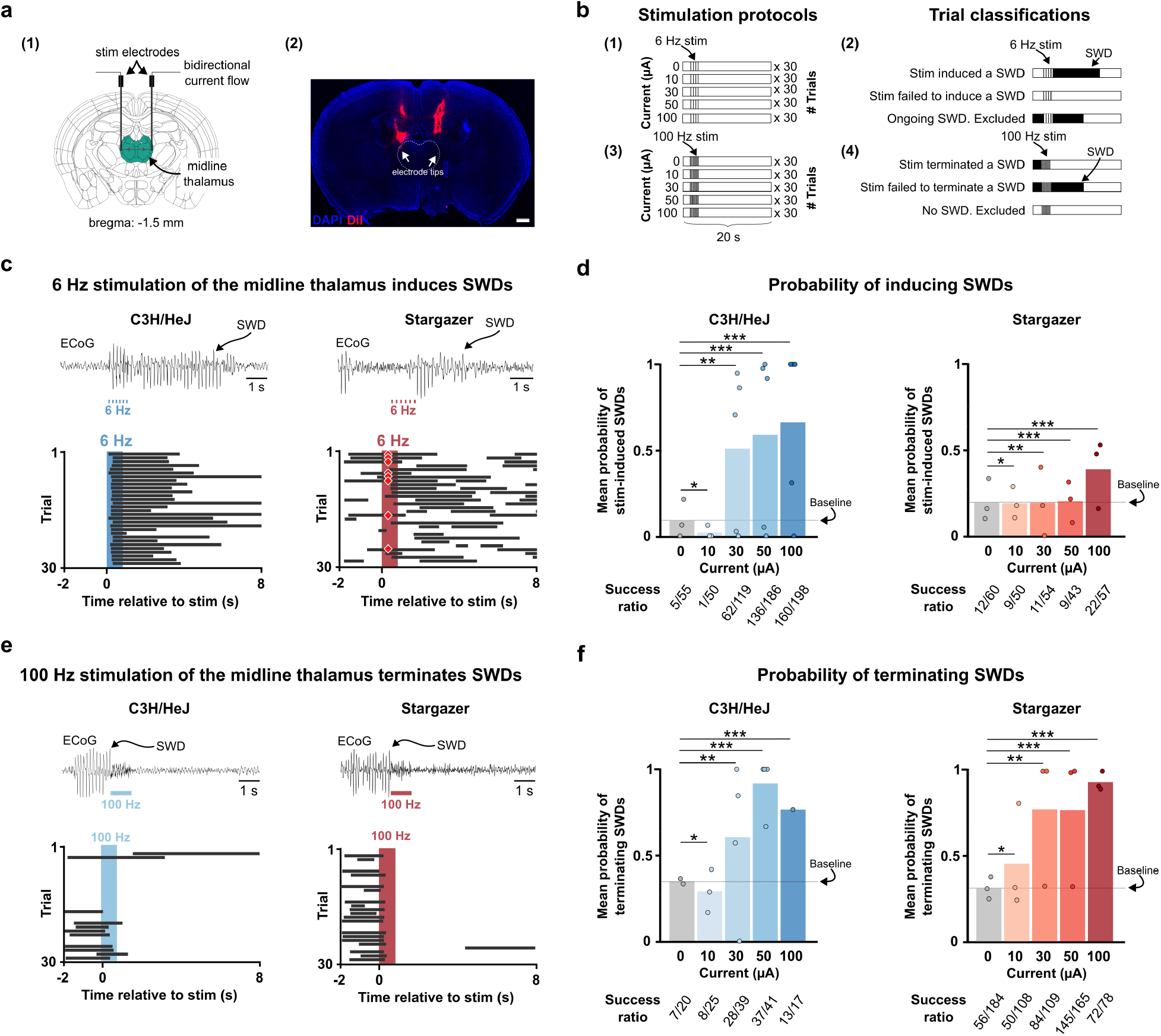
High-frequency stimulation of the midline thalamus terminates SWDs. ***a***. Approach (1) and representative brain section with visible stimulation electrodes within the midline thalamus, marked with a white dotted line (2). White arrows point to electrode tips. Scale bar: 500 µm. ***b***. Stimulation protocol for inducing (1) and terminating SWDs (3). For SWD induction, trials were classified as successful or unsuccessful based on whether stimulation induced an SWD; trials with ongoing SWDs at stimulation onset were excluded from induction analyses (2). For SWD termination, trials were considered successful if SWDs ended within the stimulation window; trials without ongoing SWDs were included in the quantification (4). ***c***. Example ECoG traces from representative trials (top) and bar graphs for 30 consecutive trials (bottom) showing reliable induction of SWDs following 6 Hz stimulation of the midline thalamus in C3H/HeJ and Stargazer mice. Red diamonds indicate trials excluded due to the ongoing SWD at the stimulation onset. ***d***. Mean probability of SWD induction as a function of delivered current for 6 Hz stimulation for C3H/HeJ (left) and Stargazer mice (right). Fractions (e.g. 5/55) below the bars show the number of successful trials over the number of all valid trials (mixed-effect model: *** p<0.001, ** p<0.01, * p<0.1. n = 5 C3H/HeJ mice, n = 3 Stargazer mice). ***e***. Example ECoG traces from representative trials (top) and bar graphs for 30 consecutive trials (bottom) showing reliable termination of SWDs following 100 Hz stimulation of the midline thalamus in C3H/HeJ and Stargazer mice. ***f***. Mean probability of SWD termination as a function of delivered current for 100 Hz stimulation, demonstrating current-dependent effect on seizure termination in both mouse models (mixed-effect model: ** p<0.001, ** p<0.01, * p<0.1. n = 5 C3H/HeJ mice, n = 3 Stargazer mice). Schematic in panel ***a*** was generated using the Scalable Brain Atlas^83^ and the Allen Institute Brain CCF Framework brain atlas^78^.

Given that spontaneous SWDs are characterized by rhythmic low-frequency activity, we hypothesized that imposing such activity within the midline thalamus would facilitate SWD generation. To test this possibility, we delivered a 1-second stimulation at 6Hz—the dominant frequency of spontaneous SWDs in C3H/HeJ and Stargazer mice—to the midline thalamus (**Fig. 6b1**). Such stimulation reliably induced self-sustained SWDs in both C3H/HeJ and Stargazer mice (**Fig. 6c**). For each tested current (i.e., 0, 10, 30, 50, and 100 µA), trials were classified as either successful or unsuccessful, depending on whether SWDs started within the stimulation window (**Fig. 6b2**). Trials with ongoing SWDs were excluded from analysis (e.g., red diamonds in **Fig. 6c**, right). The probability of SWD induction increased with stimulation current amplitude indicating a current-dependent facilitation of pathological network activity (**Fig. 6d**). However, SWD induction was far less effective in Stargazer mice, perhaps reflecting a higher baseline SWD burden in this model (see dotted lines in **Fig. 6d** and **Fig. 1f**), and an SWD refractory period that constrains the capacity for initiating additional SWDs. Nonetheless, these results demonstrate that rhythmic activation of midline thalamic nuclei at native SWD frequencies is sufficient to promote large-scale network synchrony and trigger SWDs.

We next tested whether modulating thalamic activity with high-frequency stimulation (i.e., stimulation frequencies far exceeding SWD frequency) disrupts ongoing SWDs; low-and high-frequency stimulation was performed in the same cohorts of C3H/HeJ and Stargazer mice. Since stimulation of the midline thalamus at 100 Hz has been shown to increase arousal in rodents^54^, we tested the effects of a 1-second stimulus delivered at this frequency (**Fig. 6b3**). Stimuli were delivered once every 20 seconds (i.e., open-loop stimulation). Trials were deemed successful if the SWD terminated within the stimulation window (**Fig. 6b4**). Trials without an ongoing SWD were excluded from analysis. In both mouse models, high-frequency stimulation delivered to the midline thalamus rapidly terminated ongoing SWDs in a current amplitude-dependent manner (**Fig. 6e, f**), but such stimulation was particularly effective in Stargazer mice. Nonetheless, these results demonstrate that high-frequency perturbation of thalamic activity robustly disrupts network dynamics required to sustain SWDs. Collectively, these results demonstrate that manipulating activity within midline thalamus strongly modulates SWD burden. In both mouse models, low-frequency rhythmic stimulation generally promoted SWD generation, whereas high-frequency stimulation generally suppressed ongoing SWDs. However, C3H/HeJ mice appeared more prone than Stargazer mice to stimulus-evoked SWDs, whereas Stargazer mice were more responsive to stimulation-induced termination of SWDs than C3H/HeJ mice. These findings further underscore the likelihood that SWD-generating circuits across models are different, and that high-frequency thalamic stimulation may not serve as a broadly applicable therapeutic approach.

## Discussion

Here, we aimed to resolve the spatiotemporal dynamics of neuronal activity that emerge during generalized absence seizures. To take an unbiased approach, we surveyed 26 brain structures in two different mouse models of absence epilepsy. This approach produced results consistent with recent rat studies showing that SWDs are associated with a subtle, general decrease in neuronal activity in the somatosensory cortex (S1) and ventrobasal thalamic complex (VB, somatosensory thalamus)^18^. Indeed, our results demonstrate modest changes in firing rate during SWDs in most recorded structures (**Fig. 2c**). Notably, neuronal synchrony was a far more prominent feature of SWDs than changes in firing rate, with the vast majority of structures showing some degree of coherence (**Fig. 3**) and phase-locking (**Fig. 4)** to the negative ECoG spike of the SWD. Most structures also showed increased correlated spiking activity during SWDs (**Fig. 5**). Importantly, although the spatiotemporal order of neuronal firing patterns observed during SWDs in C3H/HeJ and Stargazer mice differed—suggesting that degenerate circuit mechanisms underlie the SWD—we identified midline thalamic nuclei common to both mouse models that appear to promote network synchrony.

### A canonical thalamocortical network

A significant body of work demonstrates that thalamic structures contain neural circuits that are sufficient to support robust neuronal oscillations. Acute brain slices containing the reticular thalamus and either the lateral geniculate nucleus or the ventrobasal complex often respond to a single, brief electrical stimulus by producing seconds-long, SWD-like neuronal oscillations^7,55,56^. Such oscillations are sustained by post-inhibitory rebound burst firing among thalamocortical neurons enriched in low threshold, T-type calcium channels (i.e., T-channels, Cav3), while rhythmic inhibition from reticular thalamic neurons provides the hyperpolarization necessary to deinactivate these channels^26,44,57,58^. Indeed, ethosuximide, the first-line treatment for absence epilepsy, inhibits T-channels and suppresses burst firing, supporting the widely accepted mechanism by which the drug reduces absence seizures^59,60^. *In vivo* LFP recordings performed in rat models of absence epilepsy revealed that the onset of the SWD is consistently associated with an initial, salient signal that emerges from the somatosensory cortex before becoming apparent in the thalamus^10,11^. Collectively, these observations gave rise to the *Cortical Focus Theory*, which posits that SWDs are initiated by a bout of hyperactivity among neurons of the somatosensory cortex that subsequently recruits rhythmogenic thalamic circuits capable of sustaining an ongoing SWD^12^. Importantly, this model does not diminish the essential contribution of the thalamus but instead addresses the site of seizure initiation within the broader seizure network. Consequently, much attention has been paid to the somatosensory cortex and thalamic nuclei that reside on the lateral aspects of the brain (i.e., reticular thalamus and ventrobasal complex). In the context of SWD generation, we refer to these thalamocortical structures as *canonical*, as their contributions to SWDs have been evaluated for the past 30 years.

Questions regarding the relative contribution of the canonical thalamocortical structures to absence seizure generation emerged in 2018, when it was demonstrated that SWDs persist in the GAERS rat model of absence epilepsy despite selective T-channel blockade in the ventrobasal complex^17^. Indeed, the role of burst firing among thalamocortical neurons, a longstanding, core feature of models describing thalamic rhythmogenicity, was deemphasized, with evidence instead suggesting that *tonic* thalamocortical neuron firing induced by excitatory cortical drive supports the maintenance of SWDs. Motivated by these observations, we sought to evaluate the spatiotemporal dynamics of neuronal activity across cortical, thalamic, and subcortical structures during generalized seizures. By doing so, we find that while burst firing is not a prominent feature among thalamic neurons during the SWD, neurons within a few structures (e.g., VB, PC, PO, CA1, **Fig. S2**) appear to modestly increase their bursting activity, relative to baseline periods. Notably, burst firing among cortical neurons during the SWD is nearly identical to such firing during non-SWD periods. Thus, while T channel-mediated bursting in the cortex may be essential for SWD generation, as proposed by McCafferty et al.^17^, this firing pattern appears to represent a common mode of activity among cortical neurons and may not be specific to SWDs. Moreover, our observation that S1 lags other brain structures within a spike-wave cycle is not consistent with the hypothesis that cortical input provides an essential drive for thalamic activity during the SWD.

### Contribution of Non-Canonical Structures to SWDs

Despite extensive attention paid to the canonical structures, it is notable that early pioneers of epilepsy research proposed that diencephalic structures along the brain’s midline are critical instigators of the SWD. Decades-old experiments in cats motivated Herbert Jasper and colleagues to propose that intralaminar nuclei (i.e., MD, CM, and CL) act as “diencephalic pacemakers” that sustain SWDs^6^. Supporting this hypothesis, recordings in rats reveal prominent CL activity during SWDs^33^, whereas in humans, CM appears to mediate early propagation of generalized SWDs^14,61^. Moreover, lesioning the intralaminar nuclei in rats eliminates pharmacologically-induced SWDs^62^. Beyond their proposed role in SWDs, midline thalamic nuclei have also been implicated in seizure-associated alterations in consciousness^63^. For example, Feng et al. (2017) demonstrated that hippocampal stimulation-induced focal seizures in rats were associated with changes in CL activity, supporting a role for this nucleus in seizure-related impairment of arousal. Consistent with the hypothesis that midline thalamic structures contribute to SWD generation, we show here that (1) single neurons within such structures are engaged prior to, and during, SWDs; (2) stimulating such structures at native SWD frequencies is sufficient to initiate SWDs; and (3) stimulating such structures at high frequencies abolishes SWDs. Collectively, these results highlight the potential importance of midline thalamic structures in SWD dynamics and suggest that current models of SWD initiation and maintenance may benefit from incorporating contributions from these non-canonical structures.

### Neural Circuit Degeneracy in Epilepsy

Degeneracy is widespread in biological systems and occurs when a common functional output is produced by multiple, distinct mechanisms^64^. The possibility of neural circuit degeneracy was first formally investigated by Prinz et al.^65^, who demonstrated that similar rhythmic motor patterns generated by a three-neuron crustacean circuit can arise from a wide range of intrinsic and synaptic parameter combinations. Since then, the concept of neural circuit degeneracy has gained traction and is now considered to represent an essential component of mammalian brain function that may enhance behavioral resiliency and confer significant evolutionary advantages^66^. However, neural circuit degeneracy is also proposed to underlie several pathological conditions, including epilepsy^67,68^.

Although the concept of neural degeneracy as applied to epilepsy is in its infancy, experimental evidence supports its existence. First, a comprehensive examination of seizures observed in zebrafish, mice and humans led to the identification of electrographic features that are preserved across species^69^. Assuming that the brains of these animals are inherently different, one logical conclusion is that similar electrographic phenotypes arise from different cellular and synaptic mechanisms^70^. More recently, Codadu et al.^68^ compared seizure-like events (SLEs) in two hippocampal slice models—low-magnesium and 4-aminopyridine (4-AP)—and concluded that different epileptogenic manipulations can produce similar SLEs, even though their underlying mechanisms differ: low-Mg²⁺ SLEs are primarily driven by strong glutamatergic excitation, whereas 4-AP-induced SLEs involve strong GABAergic activity that results in neuronal depolarization. Thus, similar seizure patterns can emerge from distinct circuit-level mechanisms, underscoring the possibility that degeneracy exists in epilepsy. This conclusion may also be relevant to childhood absence epilepsy (CAE), a common polygenic form of pediatric epilepsy in which several risk genes have been identified. Notably, however, the penetrance of any single mutation within the CAE population is low, thereby giving rise to the general assumption that CAE patients carry mutations in multiple risk genes^67,72^ and that seizures can arise from multiple, distinct sets of gene mutations. If true, then the likelihood that SWDs result from degenerate circuit mechanisms is high. Although there is currently no direct evidence that SWDs in human CAE arise through degenerate circuit mechanisms, this genetic heterogeneity raises the possibility that distinct circuit perturbations may converge on a similar electrographic phenotype. Whether neural circuit degeneracy contributes to human absence epilepsy remains an important question for future studies.

Definitive evidence that degeneracy underlies SWDs is lacking, as we have not yet resolved the underlying mechanisms that generate C3H/HeJ and Stargazer SWDs. Nonetheless, from first principles, if such degeneracy exists, then it follows that: (1) the final macroscopic event—i.e., the SWD as recorded in the ECoG—must be highly similar between C3H/HeJ and Stargazer mice (**Fig. 1f**); (2) neuronal activity patterns across multiple brain structures are likely different during C3H/HeJ and Stargazer SWDs (**Fig. S3c**); and (3) SWD-generating circuits across different models likely respond differently to similar perturbations (**Fig. 6**). In this regard, our observation that the temporal relationships among active neurons across multiple structures are different between C3H/HeJ and Stargazer SWDs provides strong evidence that such highly similar seizures arise from different circuit-level mechanisms. This demonstration provides a new perspective on the challenges faced when treating epilepsy. That is, the precise targeting of specific cellular or circuit elements may have limited efficacy when multiple SWD solutions exist. Despite this conclusion, it remains possible that targeting common, model-independent circuit elements—such as our identified thalamic structures with high functional connectivity—will support a more universal treatment for absence epilepsy, albeit with variable efficacy.

### Limitations of the study

This study has limitations. Although C3H/HeJ and Stargazer mice reproduce key electrophysiological features of absence epilepsy, these models have limitations. C3H/HeJ mice exhibit retinal degeneration and blindness, whereas Stargazer mice display cerebellar dysfunction and ataxia, phenotypes that are not commonly associated with childhood absence epilepsy. Therefore, caution is warranted when extrapolating findings from these models to the human condition.

In addition, extracellular silicon probe recordings sample only a subset of neurons within each recorded brain region. As a result, neuronal populations with sparse firing, low-amplitude extracellular waveforms, or limited probe coverage may be underrepresented. Consequently, our recordings may not fully capture the diversity of neuronal activity within individual brain structures. Furthermore, cell-type classification was based on extracellular waveform features rather than genetically or molecularly defined markers. Although waveform-based approaches provide useful estimates of putative neuronal identity, they do not provide definitive cell-type classification, and some neurons may be misclassified. Thus, cell-type-specific findings should be interpreted cautiously, particularly given the limited number of classified units.

Moreover, electrical stimulation experiments demonstrate that perturbing activity within a given brain region can trigger or terminate SWDs, but they do not by themselves establish that the stimulated structure is uniquely responsible for SWD generation. Indeed, previous studies have shown that low-frequency stimulation of the somatosensory cortex can induce SWDs^73^, whereas high-frequency stimulation of several other brain regions, including the subthalamic nucleus and tuberomammillary nucleus, can suppress SWDs^74,75^. Together, these findings suggest that electrical stimulation perturbs the state of distributed seizure networks, allowing modulation of multiple interconnected nodes to shift the brain toward or away from SWD generation rather than identifying a single anatomical seizure generator. Notably, stimulation at the native SWD frequency tends to promote SWDs, whereas high-frequency stimulation suppresses them, indicating that stimulation parameters may be as important as anatomical target in determining the network response.

## Resource Availability

### Lead Contact

Further information and requests for resources and reagents should be directed to and will be fulfilled by the Lead Contact, Mark Beenhakker.

### Materials Availability

This study did not generate new unique reagents.

### Data and Code Availability

- **Data**

Spike times and ECoG from all recordings are publicly available are shared via University of Virginia Dataverse at https://doi.org/10.18130/V3/BADMP8.

- **Code**

Code used to analyze data is publicly available at https://github.com/blabuva/Thalamus-drives-2025-Dulko (DOI: 10.5281/zenodo.19392450).

- **Other items**

Any additional information required to reanalyze the data reported in this paper is available from the lead contact upon request.

## Author contributions

E.D. and M.B. designed research and experiments; E.D., S.K., A.G.C., A.R., I.L, and M.P. performed experiments; E.D., S.K., and M.B. analyzed data; E.D. and M.B. wrote the manuscript with critical review by other authors.

## Acknowledgments

This study was supported by the NIH (R01NS099586), Department of Pharmacology (University of Virginia), Ingrassia Family Echols Scholars Research Grant, a UVA Wagner Fellowship, and a UVA Double Hoo Research Grant. We thank Dr. Rajat Saxena (Kavli Institute for Systems Neuroscience) and Dr. Justin Shobe (University of California, Irvine) for sharing the headplate designs, Charlitien Long (University of California) for sharing the protocol for dye application, Dr. Jianhua Cang (University of Virginia) and Dr. Rolf Skyberg (University of Oregon) for advice regarding single unit recordings, Dr. Serapio Baca (University of Virginia) for fabricating custom 3D parts. We thank Molly Gerding (University of Virginia) for genotyping and breeding assistance. We are grateful to Dr. Daniel Meliza and Dr. Seiji Tanabe (University of Virginia) for their valuable consultation on spike-field coherence analysis, and to Dr. Marieke Jones and Clay Ford (University of Virginia) for their advice on statistical analysis. We thank the SciDraw contributors^84^ for providing the scientific illustration that was adapted for use in the graphical abstract.

## STAR Methods

## 1. EXPERIMENTAL MODEL AND STUDY PARTICIPANT DETAILS

### C3H/HeJ mouse

C3H/HeJ mice (strain #: 000659) were purchased from Jackson Laboratory (Bar Harbor, Maine). A total of 28 C3H/HeJ mice (14 males, 14 females) were used in this study. At the time of the electrophysiological recording, mice were 9.2 ± 1.2 weeks old (mean ± SD). All mice had access to food and water *ad libitum* and were kept on a 12h light:12h dark cycle. Animals were group-housed prior to surgery and singly-housed thereafter to prevent injury or damage to implants. All studies were approved by the Institutional Animal Care and Use Committee at the University of Virginia (protocol number: 3892-08-23).

### Stargazer mouse

Stargazer mice (strain #: 001756) were purchased from Jackson Laboratory (Bar Harbor, Maine). A total of Stargazer mice (6 males, 15 females) were used in this study. At the time of the electrophysiological recording, mice were 9.8 ± 2.7 weeks old (mean ± SD). Stargazer genotype was identified by standard PCR for wildtype and mutant alleles of the CACNG2 (“stargazin”) gene performed by Transnetyx (Cordova, Tennessee). All mice had access to food and water *ad libitum* and were kept on a 12h light:12h dark cycle. Animals were group-housed prior to surgery and singly-housed thereafter to prevent injury or damage to implants. All studies were approved by the Institutional Animal Care and Use Committee at the University of Virginia (protocol number: 3892-08-23).

## 2. Methods details

### Headplates and ECoG Implantation

Mice were placed in an anesthesia chamber and anesthetized with 3% isoflurane (Covertus, cat no: 029405, Portland, Maine). Next, mice were placed in a stereotaxic frame for surgical implantation of a custom-made ECoG manifold (DigiKey, Thief River Falls, Minnesota). Isoflurane was maintained at 1.5-2.5% during surgery and body temperature was maintained at 37.5°C. Carprofen was administered for pain management (2.5 mg/kg, subcutaneous). An incision was made to expose the skull, and a stainless steel headplate (Star Rapid, China) was placed on the skull and fixed in place with white C&B Metabond (Parkell, Edgewood, New York). Burr holes were drilled for two cortical electrodes (motor and somatosensory cortices, coordinates: ML:-1.0, AP: 2.0; ML:-1.5, AP:-1.5, respectively) and one reference electrode in the cerebellum (ML: +1.0, AP: +1.0). 12 C3H/HeJ mice had only one ECoG electrode (motor cortex). Electrodes were made of PFA-coated stainless steel wire (diameter 0.008 in) with stripped ends (A-M Systems, cat no: 791400, Sequim, Washington) and secured in burr holes with iBOND (Patterson Dental, cat no: 044-1113, Richmond, Virginia). Following surgery, all mice were given at least seven days of recovery.

### Electrophysiological recording

Before recording, mice were habituated to head fixation for a minimum of three hours per day for two days. On the day of the recording, mice were anesthetized with isoflurane and a 2×2 mm craniotomy was performed above the left hemisphere. The dura was removed to allow for easier probe penetration, a procedure that lasted no more than 20 minutes to minimize the influence of anesthesia on the brain. Approximately one hour later, mice were then placed on a rotatable Styrofoam ball and fixed to a headframe clamp with two screws. All neural recordings were carried out with 64-or 256-channel silicon microprobes^76^ (University of California, Los Angeles). The probe was coated with CM-DiI (Thermo Fisher, cat no: V22888, Waltham, Massachusetts) diluted 1:4 in ethanol and mounted to a 3-axis micromanipulator (New Scale Technologies, Victor, New York). The probe was manually lowered into the brain until action potentials were recorded on the electrodes closest to the probe tip. After the probe penetrated the brain to a depth of ∼300 microns, the probe was lowered automatically at a rate of 200 µm/min (total of 4 mm or less in the brain) using a micromanipulator system (New Scale Technologies, Victor, New York) to minimize damage caused by the insertion^77^. Probes were allowed to settle for 20 min after they reached their target. The craniotomy was covered with mineral oil (Sigma-Aldrich, St. Louis, Missouri) to prevent the brain from drying. Single-unit data, cortical ECoG data, and motion data were acquired simultaneously at 30 kHz with the RHD recording system (Intan Technologies, Los Angeles, California) for 30 - 60 minutes. All recordings were made in awake, head-fixed mice and were obtained between 11 am and 2 pm. During this period, an average of 0.38 seizures/min (C3H/HeJ) and 1.21 seizures/min (Stargazer) were recorded.

### Histology

Histological analyses were performed to confirm the position of the silicon probe in brain tissue from all mice. Shortly after the end of each recording, mice were perfused with 1x PBS followed by 4% PFA. After an additional post-fixation period of two days in 4% PFA at 4°C, the brain was sectioned into 70 µm sections. Brain sections were stained for parvalbumin protein using free-floating immunofluorescence to visualize the reticular thalamus (for anatomical reference). Sections were placed in 0.1% sodium borohydride (Millipore Sigma, cat no: 452882, Burlington, Massachusetts) in PBS for 15 minutes before being washed twice in 1x PBS. Sections were then incubated in a 3% normal donkey serum (Abcam, cat no: ab7475, Cambridge, United Kingdom) in 0.5% Triton X-100 (Millipore Sigma, cat no: T8787) blocking solution overnight at 4°C. Next, sections were washed in 1x PBS three times before being incubated in monoclonal anti-parvalbumin antibody produced in mouse (Millipore Sigma, cat no: P3088), 1:2000 dilution in 1% normal donkey serum (Abcam, cat no. ab7475, Waltham, Massachusetts) at 4 °C overnight. Sections were then washed in 1x PBS four times and incubated in Alexa Fluor® 488 AffiniPure Donkey Anti-Mouse IgG (H+L, Jackson ImmunoResearch, cat no: 715-545-150, West Grove, Pennsylvania), 1:200 dilution in 1% normal donkey serum in 0.5% Triton X-100 overnight at 4°C. Finally, sections were washed in 1x PBS four times before being mounted to Shandon Colorfrost™ Plus microscope slides (Thermo Fisher Scientific, cat no: 99-910-01, Waltham, Massachusetts). After allowing slides to dry, sections were rinsed with deionized water and stained for cell nuclei using DAPI Fluoromount-G® (SouthernBiotech, cat no: 0100-20, Birmingham, Alabama) and covered using a 24×50 mm No. 1.5 VWR coverslip (VWR, cat no: 48393-081, Bridgeport, New Jersey). Slides were imaged using a Zeiss Imager.Z1 and Zeiss AxioCamMRC and Zeiss Objective N-Achroplan 5x/0.13 Pol M27 (WD = 5.5 mm). The X-Cite XYLIS Broad Spectrum LED illumination system (Excelitas Technologies, Waltham, Massachusetts) was used for excitation. Fluorescence from DAPI, DiI, and Alexa Fluor 488 was collected using the DAPI, Cy3, and YFP channels, respectively, using Neurolucida software (MBF Bioscience, Williston, Vermont). Images with visible probe tracks were downsized using ImageJ (National Institutes of Health, Bethesda, Maryland). Probe trajectories were reconstructed by aligning images with the Allen CCF^78^ atlas using publicly available MATLAB (MathWorks, Natick, Massachusetts) code (https://github.com/petersaj/AP_histology)^79^.

### Spike Sorting

Neural signals acquired from silicon probes contained either 64 or 256 recording channels. Offline spike detection and automatic sorting was performed with Kilosort 2.5^80^ and detected clusters were manually curated with Phy^81^. Clusters with less than 500 spikes were not considered. Clusters with more than 500 spikes were considered for single-and multi-units. Only clusters associated with a clean and consistent waveform shape, a normal distribution in the amplitude view, and auto-correlograms showing few to no spikes falling within the refractory period were classified as single-units. Units were classified as multi-unit if more than 20% of spikes in the amplitude view were cut off, or the cross-correlogram suggested no refractory period. After manual curation, resultant single-and multi-unit clusters were imported into MATLAB for further analysis using custom code.

### Unit classification

Recorded single units were classified as putative excitatory or putative inhibitory neurons based on two extracellular spike waveform features: trough to peak time and half-amplitude duration^41^. Trough to peak time was measured from the electrode channel with the largest spike amplitude by identifying the minimum voltage value (trough) of the mean spike waveform and the subsequent maximum (peak); the time interval between these two points was defined as the trough to peak time. Half-amplitude duration was calculated as the time between the initial crossing of one-half of the trough amplitude and the subsequent return to that value after the trough (**Fig. S1e**). Distribution of classified units and representative waveforms are shown in **Fig, S1f**. Units with a half-amplitude duration > 0.3 ms were classified as putative excitatory. Units with a half-amplitude duration < 0.3 ms and a trough to peak duration < 0.5 ms were classified as putative inhibitory. Units that did not satisfy either criterion were classified as unclassified and were excluded from cell-type analyses.

### Tonic and Bursting Classification

Instantaneous firing rates (IFRs) were calculated for each single unit during both non-seizure and SWD periods. For each brain structure, spike times from all single units were extracted to compute IFRs. IFRs below 200 Hz were classified as tonic, whereas IFRs at or above 200 Hz were classified as burst firing. The total number of IFRs within a given brain structure was considered 100%, and the percentages of tonic and burst firing were calculated relative to this total (**Fig. S2a, b, f**). Inter-burst intervals were computed by first identifying bursts, defined as sequences of three or more spikes with an inter-spike-interval (ISI) of ≤ 7ms that were preceded by a silent period ≥ 100 ms^17^ (**Fig. S2c-e**). After bursts were identified, the duration between two consecutive bursts was measured to calculate the inter-burst interval.

### Seizure Detection Algorithm

ECoG signals recorded from motor and somatosensory cortex were imported into MATLAB. Candidate seizure events were detected by first computing a spectrogram with 1 second windows and 75% window overlap. Power between 4-8 Hz was subsequently calculated for all windows and a threshold equal to the 95^th^ percentile of this distribution was used to identify candidate events in the ECoG trace. Events briefer than 500 ms were excluded and events separated by less than 1 second were merged. Negative peaks (troughs) during these events were identified and only those greater than 3 standard deviations from the overall EEG voltage distribution were retained. If two negative peaks were detected within 50 ms, the second timepoint was excluded. These candidate events were then manually verified by an experienced scorer. Seizure onset was defined as the emergence of the first SWD-associated negative peak in the motor cortex ECoG that exceeded 3 standard deviations from the overall ECoG voltage distribution.

### Stimulation Experiments

Electrical stimulation experiments were performed in awake, head-fixed C3H/HeJ and Stargazer mice. Stimulation electrodes were made by twisting PFA-coated stainless steel wire (diameter 0.002 in, A-M Systems, Sequim, Washington) and soldering them to female pins (DigiKey, Thief River Falls, Minnesota). Electrodes targeting midline thalamus were implanted bilaterally using the following coordinates: AP:-1.5 mm, ML: 0.7 mm, DV: 2.7 mm. Prior to insertion, electrodes were coated with CM-DiI (Thermo Fisher, cat no: V22888, Waltham, Massachusetts) diluted 1:4 in ethanol. For these experiments, the ECoG electrode was implanted in the pre-frontal cortex to allow for SWD detection. Mice were given a week of recovery between implantation and experimentation. To quantify the probability of inducing and terminating SWDs, we employed an automated stimulation protocol where the stimulation was triggered every 20 seconds for 1 second (1 sec ON, 19 seconds OFF; current: 0, 10, 30, 50, 100; frequency: 6 or 100 Hz; pulse width: 1ms). Mice underwent three recording sessions conducted on separate days. SWD onset and offset were labeled manually after the experiment. Mice with fewer than five valid trials per condition (e.g., 0 µA current, 6 Hz frequency) were excluded from the analysis.

## 3. QUANTIFICATION AND STATISTICAL ANALYSIS

### Data Analysis

All data analysis was performed using custom-written MATLAB scripts as well as publicly available toolboxes: FMAToolbox (https://github.com/michael-zugaro/FMAToolbox) and Chronux (https://chronux.org/).

### Spike-Field Coherence

Spike-field coherence was computed to evaluate the relationship between neuronal firing in each recorded brain structure to the ECoG signal (**Fig. 3** and **Fig. S3b**). Coherence is a measure of phase consistency between two time-varying signals and values range between 0 and 1, wherein 0 indicates no coherence and 1 indicates strong coherence. Power and coherence spectra were estimated using Welch’s method with a Hamming window of 500 ms and 50% overlap, applied to both spike counts and ECoG. Power spectra for ECoG were computed using the Fast Fourier Transform (FFT) with 1024 points and a sampling rate of 1 kHz (Sxx, see equation below). Power spectra for firing activity were estimated from binned spike counts (Syy). Bin size was adjusted to match the length of the seizure. Seizures briefer than 0.5 seconds were excluded from the analysis. Cross-spectral densities were estimated by multiplying complex Fourier coefficients of spike counts and LFP signals, and then averaging across time windows (Sxy).

Next, coherence for each seizure was computed using the equation below and then averaged across all seizures.

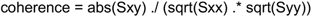

### Phase analysis

Action potential times were defined relative to the large, negative-going ECoG spike of the SWD (**Fig. 4a-d** and **Fig. S3b, c**). First, a single cycle of the SWD was defined as the duration between the peak of one negative ECoG spike and the peak of the subsequent, negative ECoG spike. Then, action potential times were expressed relative to a normalized SWD cycle (i.e., phase), wherein 0 and 1 correspond to the start and end of the SWD cycle, respectively. Phase values were summed across seizures and animals on a per structure basis and then presented in histogram format (bin size: 0.01 phase). For analyses of putative excitatory and inhibitory neurons, spike times were analyzed separately for each cell type. Phase locking was quantified using vector analysis, in which vector strength measured the concentration of spikes around the preferred firing phase (higher values indicate stronger phase locking), and the preferred phase was defined as the center of mass of the phase distribution. In **Fig. 4c** (right), dot position indicates the preferred phase, and dot size is proportional to vector strength (phase-locking strength).

### Entire Spiking Activity (ESA)

ESA was computed for all electrodes in all recordings with the following algorithm: (1) bandpass filter the original 30kHz-sampled voltages between 0.3-12.5 kHz offline using a FIR filter; (2) rectify the filtered signal; (3) convolve the rectified signal with a Gaussian kernel (σ = 5ms) evaluated over 201ms; and (4) downsample the convolved signal to 3kHz (**Fig. S5a**). The algorithm generates a continuous ESA vector with a final sampling rate of 3kHz. In some cases, as in the analysis of temporal order across brain structures, ESA vectors were z-scored after downsampling. ESA proved particularly useful for evaluating neuronal rhythmicity in brain structures wherein a limited number of single units was isolated, such as RT and ethmoid nucleus of thalamus (EtH), because peaks in cross-correlational analyses are generally only revealed with a high sample size.

Oscillation indices (OIs) of ESA autocorrelations were computed to evaluate spiking rhythmicity during SWD and non-seizure periods. First, every sample in the ESA vectors was labeled as “SWD” if it occurred during a SWD or “non-seizure” otherwise. Then, autocorrelations were calculated for the SWD and non-seizure periods and normalized such that their identity peaks at t_0_ were equal to 1. Finally, OI was calculated by subtracting the value of the first trough off the identity peak (Autocorr_trough_) from the value of the first non-identity peak (Autocorr_peak_) of the autocorrelation (i.e., Autocorr_peak_ - Autocorr_trough_) (**Fig. S5b-f**). If no non-identity peaks were present, the OI was set to 0. OI values range from 0 to 1. Finally, to resolve how neuronal rhythmicity differs between non-seizure and SWD epochs, the difference between respective OIs was calculated (i.e., OI_SWD_ - OI_non-SWD_). Therefore, the final value can range from-1 to 1.

To quantify the magnitude of phase-locking and phase preferences of ESA during SWD, mean ESA vectors relative to SWD phase were computed. To this end, first, all SWD cycles were divided equally into 100 phase bins between 0 and 1, with 0 corresponding to the negative peak of SWD (**Fig. S8a**). Then, the mean ESA in each bin was calculated such that each SWD cycle had an ESA value for all 100 phase bins. The average across all SWD cycles was then taken; these are the ESA-SWD phase distributions (**Fig. S8b**). Finally, mean vectors of these distributions were computed. Mean vector lengths (MVL) correspond to the magnitude of ESA-SWD phase-locking. Mean vector angles (θ) correspond to the ESA-SWD phase preference (**Fig. S8**).

For all ESA analyses described above, electrodes that did not have at least one well isolated single unit, as determined by manual curation following spike sorting, were excluded. The reason for this is to ensure that electrodes not recording any spiking activity were not included in analysis; such electrodes would artifactually decrease the measures of ESA-SWD relationship because of insufficient sampling of neural activity.

### Synchrony Analysis

Intra-nucleus comparisons: spike times of all neurons within each brain structure were binned (25 ms, bin size) and used as an input for pair-wise Pearson correlation analysis^51^ (**Fig. 5b** and **Fig. S3e**). Brain structures with fewer than five neurons were excluded from this analysis.

Inter-nucleus comparisons: spike times of all neurons from different brain structures (but recorded within the same recording session) were binned (50 ms, bin size) and used as an input for pair-wise Pearson correlation analysis^51^ (**Figure 5c, d**). A larger bin size was chosen to capture delayed correlations between distant brain regions, which might be missed with a smaller bin size. Threshold values were selected empirically to retain a consistent proportion of high correlations between mouse models. Specifically, a threshold of 0.15 was used for C3H/HeJ mice, retaining the top 21% of correlations, while a threshold of 0.1 was used for Stargazer mice, corresponding to the top 6% of correlations.

### Lag Analysis

Spike times during SWDs were binned (5 ms, bin size) and used for cross-correlation analysis to determine the temporal relationship between distinct brain structures (**Fig. 4g, Fig. S3d,** and **Fig. S6**). Only neurons recorded in the same recording session were analyzed. Next, center of mass of the cross-correlation distribution was calculated within ± 50 ms lag. The position of this center was taken as the estimated lag (positive values indicating that the compared structure lagged the reference, negative values indicating that it led).

## Statistical Analysis

Statistical significance of SFC was tested in MATLAB using a nonparametric surrogate approach (**Fig. 3** and **Fig. S3a**). Specifically, 200 surrogate spike count series were generated by randomizing the observed values to disrupt temporal alignment of neuronal firing, while preserving the average firing rate. This ensures that any change in coherence is not due to a change in the average firing rate, which is known to influence coherence measures. Then, the mean of observed coherence within the 4.5–6.5 Hz band was compared against the empirical surrogate distribution. If the observed value was higher than the 95^th^ percentile of the surrogate distribution, the coherence was considered statistically significant.

To assess whether phase distributions differed between SWD and non-seizure epochs, we imposed a surrogate oscillation of matched SWD frequency during non-seizure epochs to provide a comparable phase reference (**Fig. 4c** and **Fig. S3b**). Spike time counts were then expressed as a function of phase for both conditions. Distributions were compared using the non-parametric Wilcoxon rank-sum test in MATLAB.

To assess differences in synchrony across brain structures (i.e., intra-nucleus correlations), we fit a linear mixed-effects model (**Fig. 5b** and **Fig. S3e**). Mixed-effects modeling was performed using the lme4 R package^82^. Each neuronal pair was treated as a datapoint. Mouse identity was included as a random intercept to account for repeated measurements across structures within the same animal. Post hoc pairwise comparisons were performed with p-values adjusted for multiple comparisons.

To assess the effect of stimulation current on SWD induction and termination probability, we fit a linear mixed-effects model using the lme4 R package^82^ (**Fig. 6**). Trial outcome (success vs. failure) was treated as a binary dependent variable, and stimulation current was included as a fixed effect. Mouse identity was included as a random intercept to account for repeated measurements within animals. Post hoc comparisons between each stimulation current and the baseline (0 µA) were performed with p-values adjusted for multiple comparisons.

Remaining statistical tests can be found in figure legends.

## Commonly used abbreviations

BG: basal ganglia
CA1, 2, 3: hippocampal CA1, 2, 3
CL: centrolateral thalamus
CM: central medial thalamus
CP: caudate putamen
DG: dentate gyrus
EtH: ethmoid nucleus of thalamus
HF: hippocampal formation
HY: hypothalamus
IMD: intermediodorsal thalamus
LD: lateral dorsal thalamus
LGN: lateral geniculate nucleus of thalamus
LH: lateral habenula
LP: lateral posterior nucleus of thalamus
M: motor cortex
MD: mediodorsal thalamus
MH: medial habenula
PC: paracentral nucleus of thalamus
PO: posterior complex of thalamus
PV: paraventricular thalamus
RT: reticular thalamus
S1: primary somatosensory cortex
SWD: spike-wave discharge
SU: subthalamic nucleus
VB: ventrobasal complex of thalamus
VPM: ventral posteromedial nucleus of thalamus
VPL: ventral posterolateral nucleus of thalamus.

