## Supplementary material for "Midline Thalamic Neurons Show Early Hypersynchrony During Absence Seizures in Mouse Models"

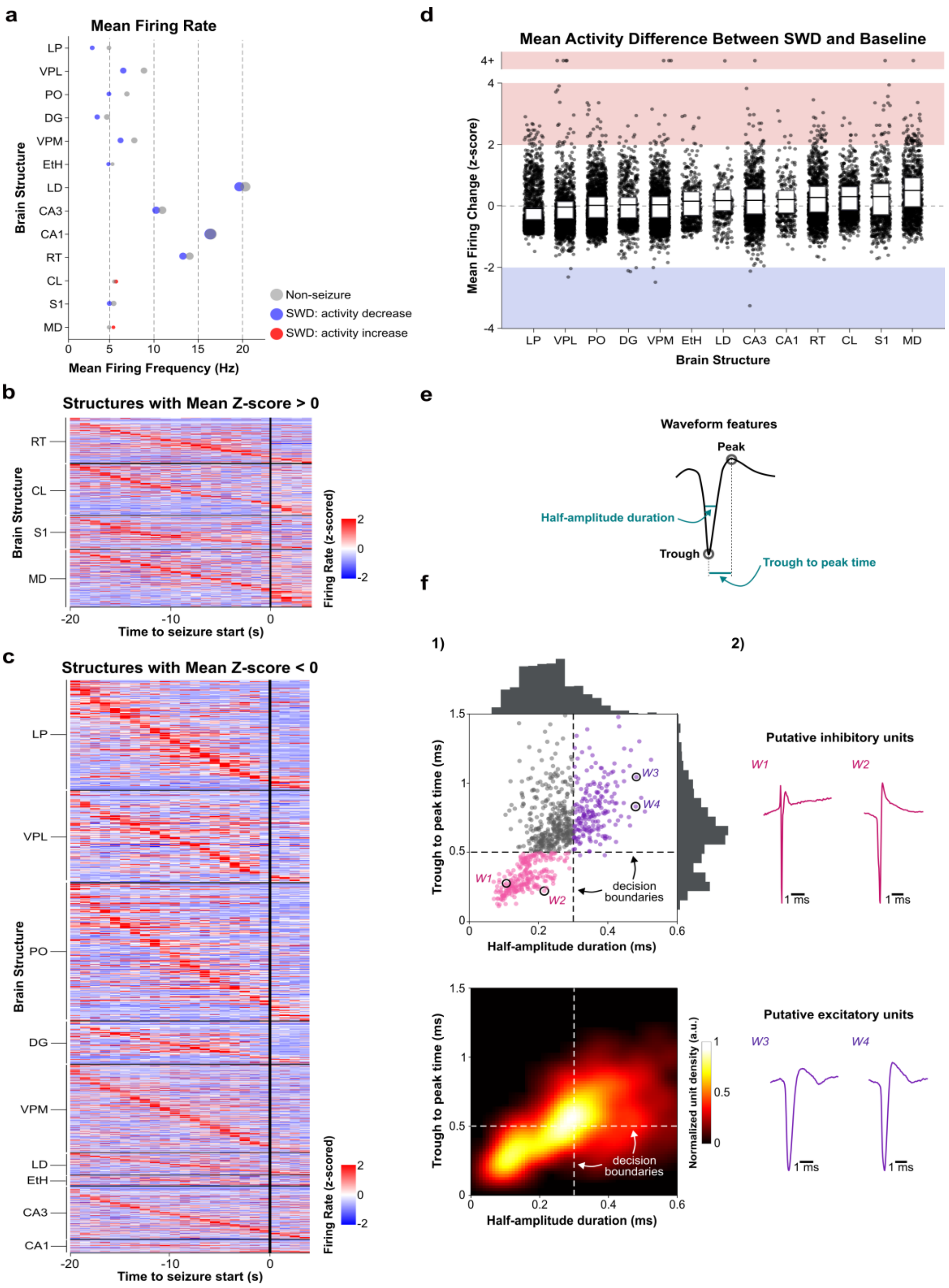


**Figure S1. SWDs produce only subtle changes in the mean firing rate in Stargazer mouse neurons**, Related to Figure 2. ***a.*** Dot plot showing mean firing rate for non-seizures and SWDs across all brain structures. Dot size is scaled based on standard deviation. Blue and red dots correspond to structures wherein activity levels trended lower or higher, respectively, during the SWD versus non-seizure periods. ***b***. Brain structures with a positive mean z-score. ***c***. Brain structures with negative mean z-score. ***d***. Box plot of mean firing change for each brain structure. Each dot represents one neuron during one seizure. The bottom and top of each box correspond to the $1^{st}$ (Q1, $25^{th}$ percentile) and $3^{rd}$ quartile (Q3, $75^{th}$ percentile), respectively; the line inside the box indicates the mean. Blue and red rectangles denote z-score < -2 and >2 respectively. ***e.*** Unit classification based on extracellular waveform features. Schematic illustrating the waveform features used for classification: half-amplitude duration and trough to peak time. ***f.*** Scatter plot of half-amplitude duration versus trough to peak time for all recorded Stargazer units. Circled points correspond to the representative inhibitory (left) and excitatory (right) waveforms shown next to the scatter (2). Heatmap shows the normalized density of units across waveform feature space. n = 21 Stargazer mice.

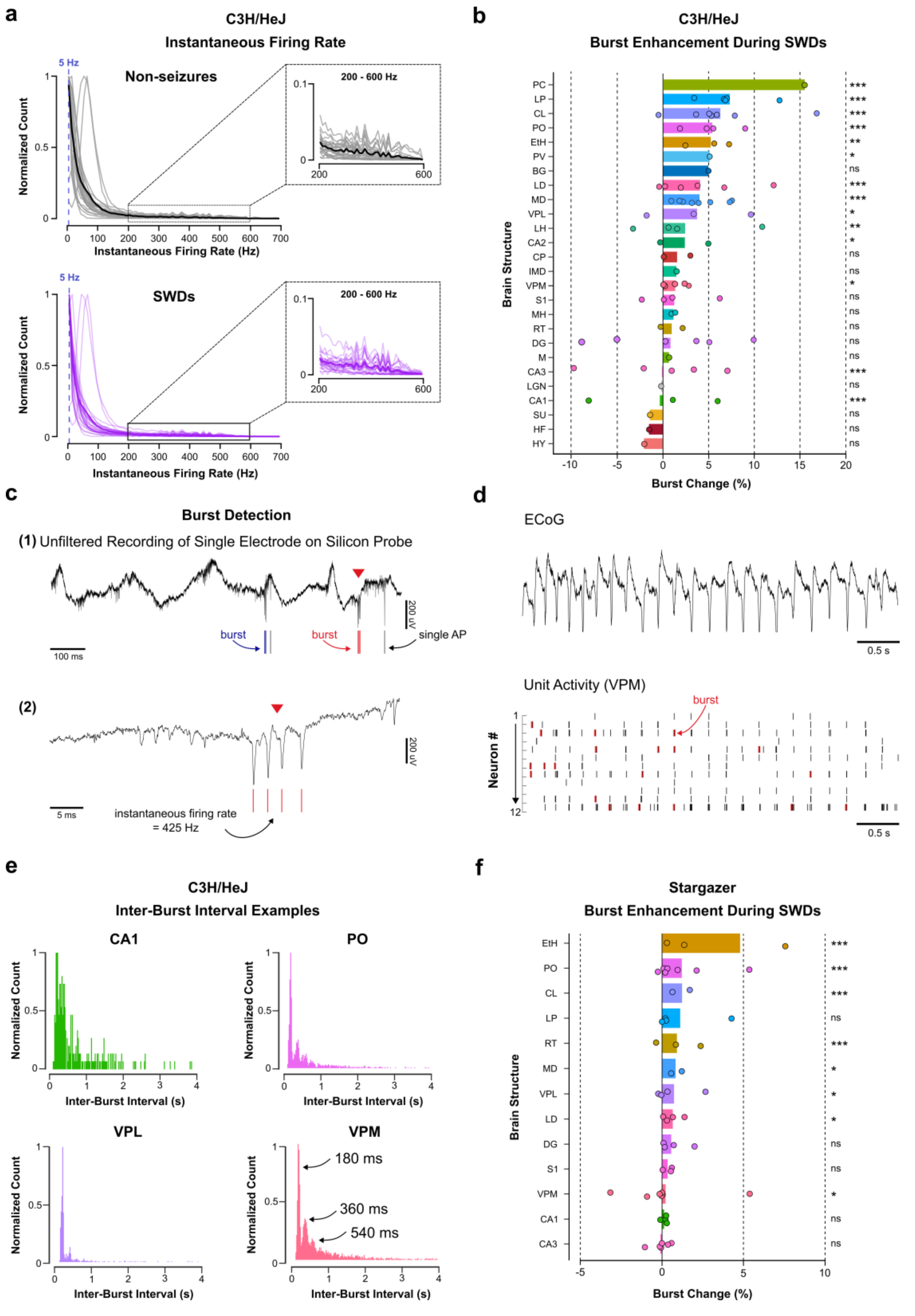


**Figure S2. Tonic and bursting firing modes during SWD in C3H/HeJ and Stargazer mice,** Related to Figure 2. ***a.*** Normalized count of instantaneous firing rate during non-seizures (top) and seizures (bottom) for all brain structures. Black and purple thick lines represent means. Skewed instantaneous firing rate is observed during non-seizures and SWDs, with the peak in 0-10 Hz range (5.8 Hz, blue dashed line). A modest level of burst firing (as defined by neurons with instantaneous firing rates greater than 200 Hz) is observed in both conditions as seen in insets (200 – 600 Hz). Histogram bin size: 10 Hz. ***b***. Bar graph showing increase in burst firing during SWDs compared to baseline. Dots represent different mice. N = 28 C3H/HeJ mice. ***c***. Representative bursts visible in the raw voltage recorded on one of the silicon probe electrodes. (1) Raw voltage trace from one of the channels on the silicon probe. Red arrow points to the second of two bursts. Below is a raster plot of action potentials of the corresponding single unit in VPM. (2) Closer look at the second burst. Despite different amplitudes of action potentials in one burst, all action potentials are assigned to a single neuron. ***d***. LFP (cortex) and raster plot of firing activity in VPM. Red lines mark detected bursts. Bursts were defined as sequences of three or more spikes with an inter-spike-interval (ISI) of ≤ 7ms preceded by a silent period ≥ 100 ms[14]. ***e***. Histograms (20 ms, bin size) of inter-burst intervals for selected C3H/HeJ brain structures for which bursting increased (based on panel b). If neurons were firing every cycle of the SWD, we would expect a single peak at 180 ms. However, the presence of peaks at 180, 360, and 540 ms indicates that many bursts occur every few SWD cycles. ***f***. Bar graph showing increase in burst firing during SWDs compared to baseline. Dots represent different mice. N = 21 Stargazer mice. Statistical comparisons in panels ***b*** and ***f*** were made using linear mixed models to account for mouse-specific effects. Asterisks represent statistical significance in pairwise post hoc two-sided comparisons: *** p<0.001, ** p<0.01, * p<0.05. (2).

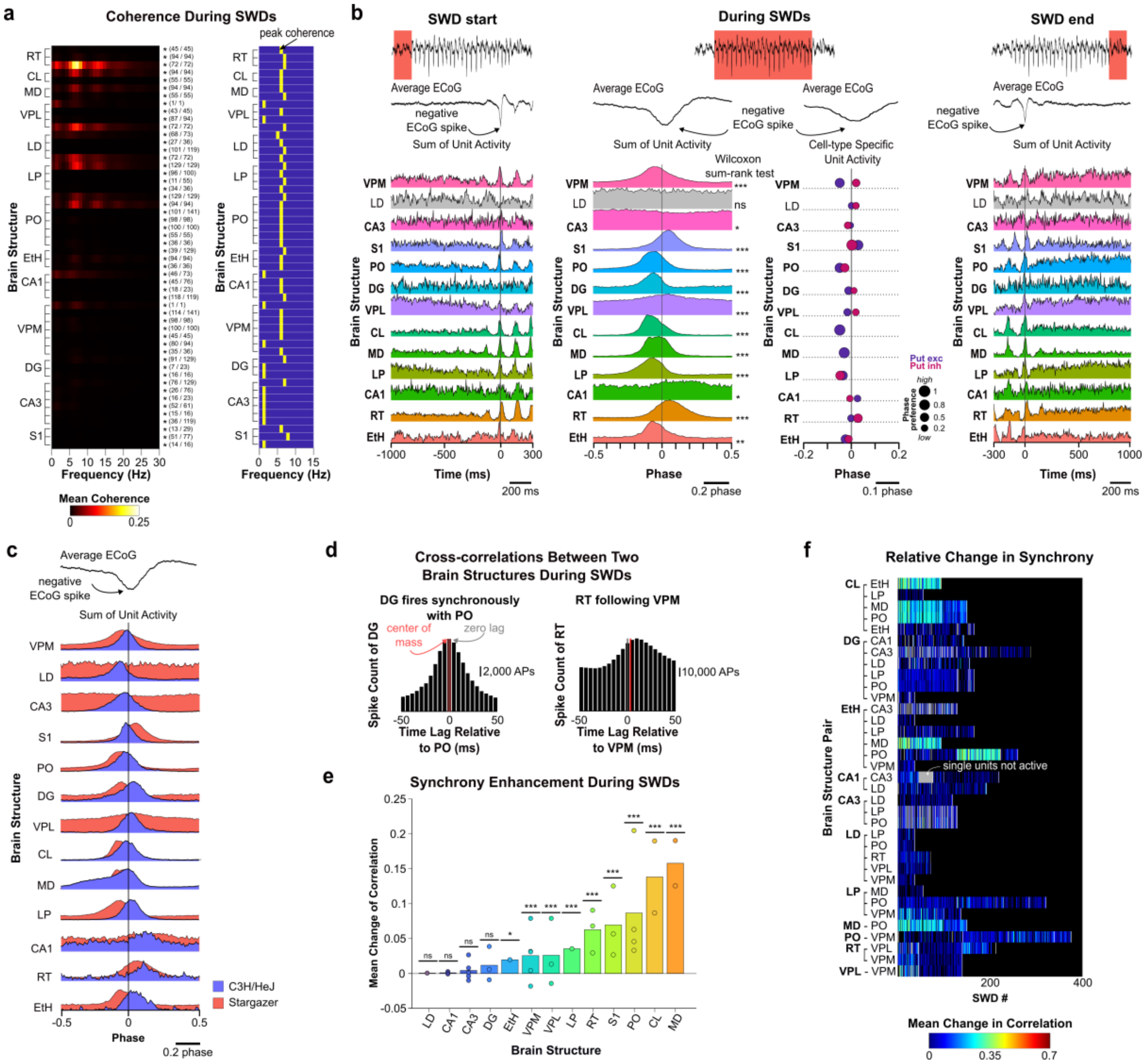


**Figure S3. Synchrony analysis for Stargazer mice**, Related to Figures 3, 4, and 5. ***a.*** Heatmap of spike-field coherence (SFC) during seizures (left). Asterisks (*) mark rows where SFC in the 4.5-6.5 Hz band was statistically significant. Fractions (e.g., 45/45) show the number of statistically significant SWDs over the total number of SWDs (Surrogate-based significance test, * $p < 0.05$). A complementary heatmap of peak SFC values highlights a consistent maximum at 5 Hz (right). ***b***. Histograms of firing for all brain structures before, during, and at the end of SWDs. Data from all mice are represented in a single histogram. Spike count was normalized to the peak firing, so peak height is the same across all brain structures. Histograms are ordered based on firing order for C3H/HeJ mice (see Fig. 4b). Scatter plot shows phase preference for putative excitatory and inhibitory neurons; dot size indicates the magnitude of phase-locking. ***c***. Difference in phase between C3H/HeJ and Stargazer mice during SWDs. n = 21 Stargazer mice, n = 28 C3H/HeJ. ***d***. Cross-correlation analysis reveals the order of firing during SWDs (e.g., RT follows VPM, which is consistent with the firing order in panel b). ***e***. Bar graph showing increase of synchrony during SWDs compared to baseline. Bars are sorted according to the mean synchrony increase. Dots represent different mice. Statistical comparisons were made using linear mixed models to account for mouse-specific effects. Asterisks represent statistical significance in pairwise post hoc two-sided comparisons: *** p<0.001, ** p<0.01, * p<0.05. ***f***. Heatmap showing the mean increase of synchrony compared to baseline (time-matched non-seizure event). Each line represents an average correlation increase for all neurons during one SWD. White bars separate seizures from different mice. Gray bars indicate that single units were not active during this seizure. n = 21 Stargazer mice.

**a**

**C3H/HeJ mouse**

| Threshold | 0.025 | 0.05 | 0.1 | 0.15 * | 0.2 | 0.25 |
|---|---|---|---|---|---|---|
| Correlations retained | 75% | 56% | 37% | 21% | 12% | 6% |

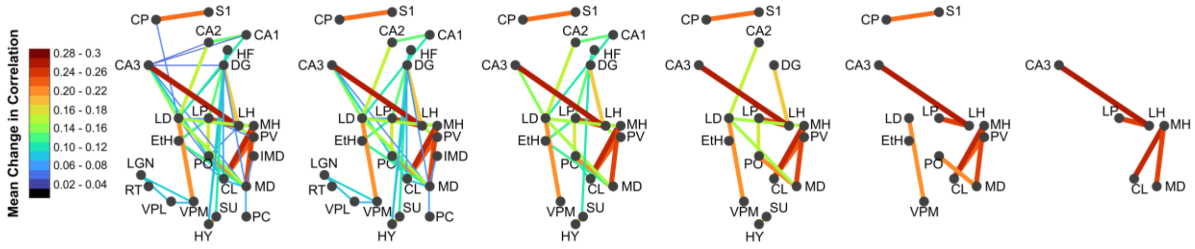


**b**

**Stargazer mouse**

| Threshold | 0.025 | 0.05 | 0.1 * | 0.15 | 0.2 | 0.25 |
|---|---|---|---|---|---|---|
| Correlations retained | 18% | 8% | 6% | 5% | 2% | 2% |

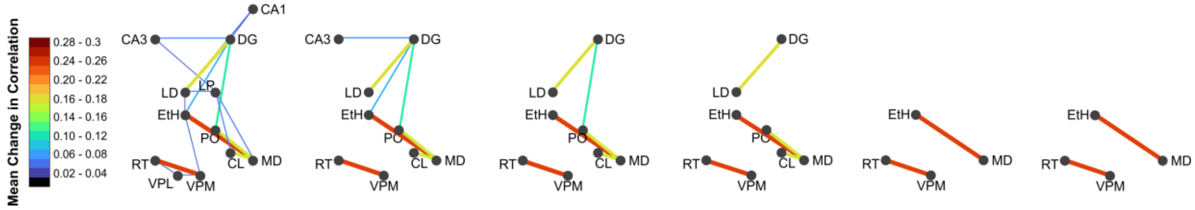


**Figure S4. Effect of correlation threshold on functional connectivity graphs**, Related to Figure 5. Functional connectivity graphs derived from Pearson correlations for C3H/HeJ (***a***) and Stargazer (***b***) mice across a range of correlation thresholds. Circles represent brain structures positioned according to their approximate anatomical location. Edges represent functional associations exceeding the indicated correlation threshold. The percentage of retained correlations is shown above each graph. Red asterisks indicate the threshold used in the main analysis (**Fig. 5d**).

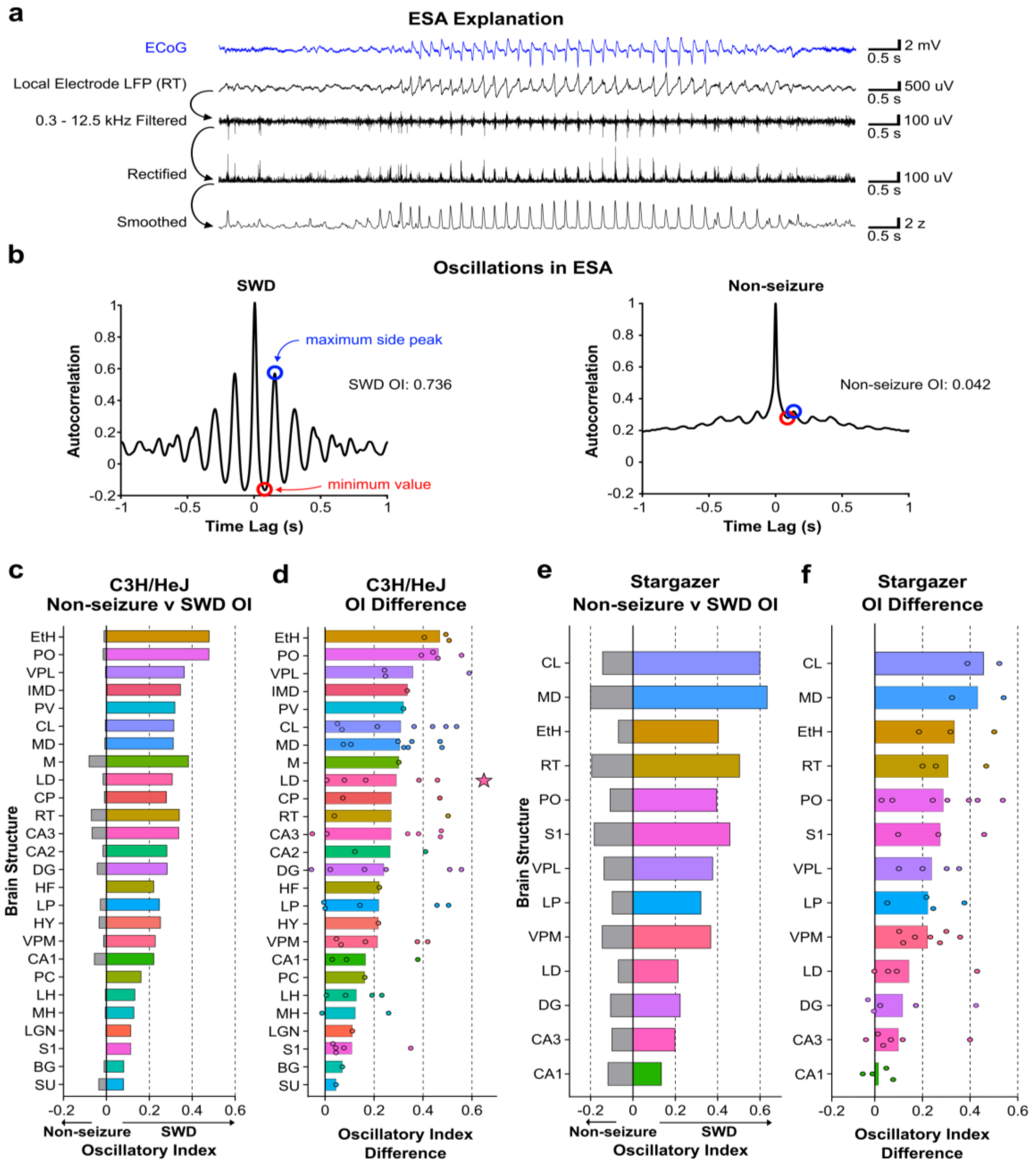


**Figure S5. Entire Spiking Activity (ESA) reveals oscillatory neuronal activity during SWDs**, Related to Figure 3. ***a***. Top trace in blue: LFP from an epidural electrode at the cortical surface used to identify the start and end of each SWD, as well as the times of negative peaks (maximum negative values of LFP spikes). Lower traces in black: steps used to compute ESA from the original raw LFP taken from one example depth electrode in RT. First, the raw data from a single channel is bandpass filtered between 0.3 and 12.5 kHz. Then, the filtered signal is rectified by taking the absolute value of the trace. Finally, the signal is convolved with a Gaussian kernel and, for some applications, the trace is transformed into a vector of z-scores. Only ESA from channels with at least 1 well-isolated neuron was used to ensure that channels without spiking activity were excluded. ***b***. Example autocorrelograms from ESA on a single electrode in LD. The SWD autocorrelation is computed on the ESA signal during the SWD and the non-seizure autocorrelation is computed from ESA outside the SWD. The oscillation index (OI) is calculated by finding the maximum side peak (blue circle) and subtracting from it the minimum value (red circle) between that side peak and the identity peak at t = 0. The OI difference is then computed by subtracting the non-seizure OI from the SWD OI. Positive OI difference scores then indicate increases in rhythmic activity during SWD. n = 1 C3H/HeJ mouse. ***c***. Average non-seizure and SWD OI for various brain regions recorded. Bars to the left in gray indicate the average non-seizure OI while bars to the right in color indicate the average SWD OI of a given brain region. ***d***. OI differences across brain structures. Circles show the average OI difference for a given brain region in a single mouse. Bars represent the mean across mice. The larger star marker in the lateral dorsal thalamus is the value computed from the example autocorrelograms in panel b. ***e.*** Average non-seizure and SWD OI for Stargazer mice. ***f***, OI differences for Stargazer mice.

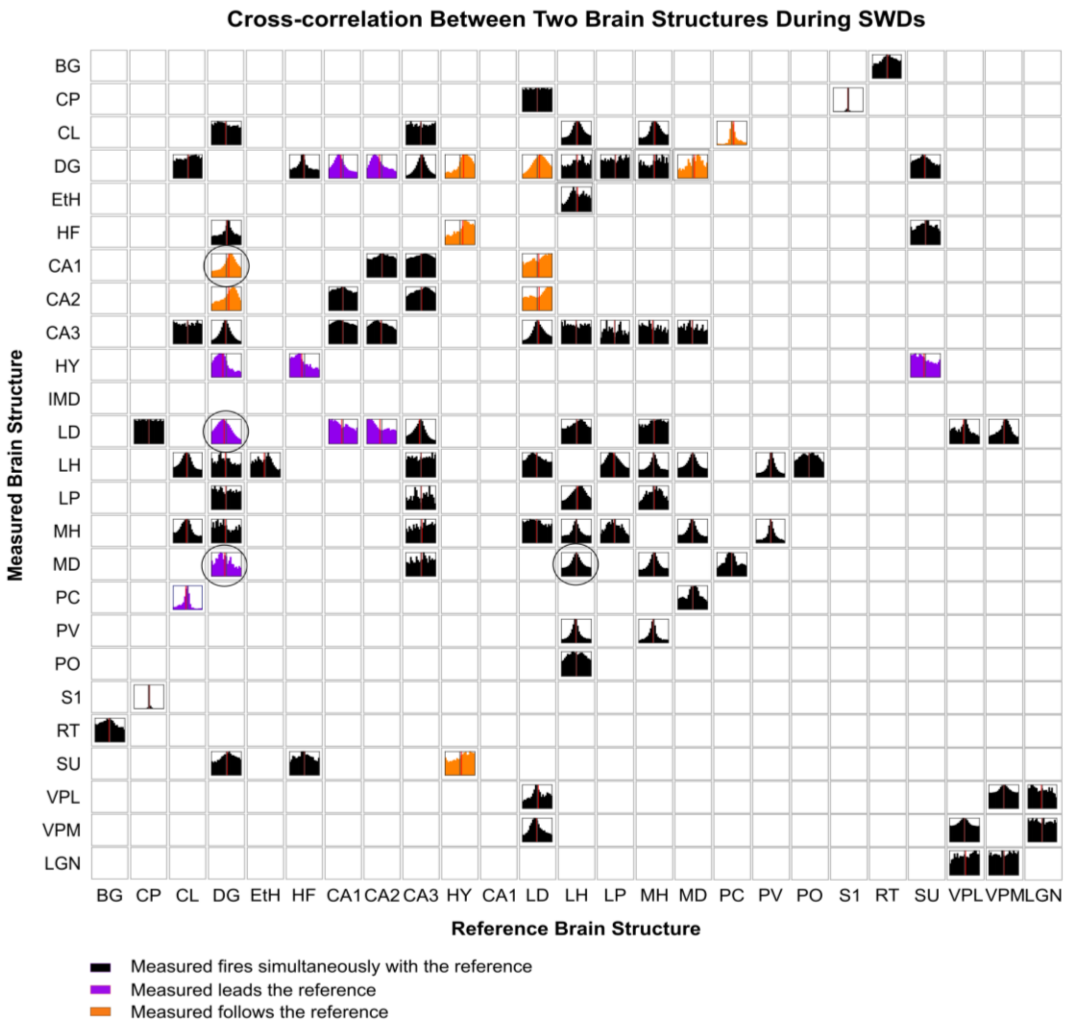


**Figure S6. Temporal firing order during SWDs in C3H/HeJ mice**, Related to Figure 4. Each histogram shows the cross-correlation between all pairs of neurons from two distinct brain structures, with columns representing reference structures and rows representing measured structures. The gray vertical line marks zero lag, and the red line indicates the center of mass (average lag) of each histogram. Histograms are color-coded based on the center of mass: black if between –5 and 5 ms (no clear lead/lag), purple if < –5 ms (measured structure leads the reference), and orange if > 5 ms (measured structure follows the reference). Cross-correlations are shown over a ±50 ms window with 5 ms bin size. Purple circles mark selected comparisons shown in Figure 4g. n = 28 C3H/HeJ mice.

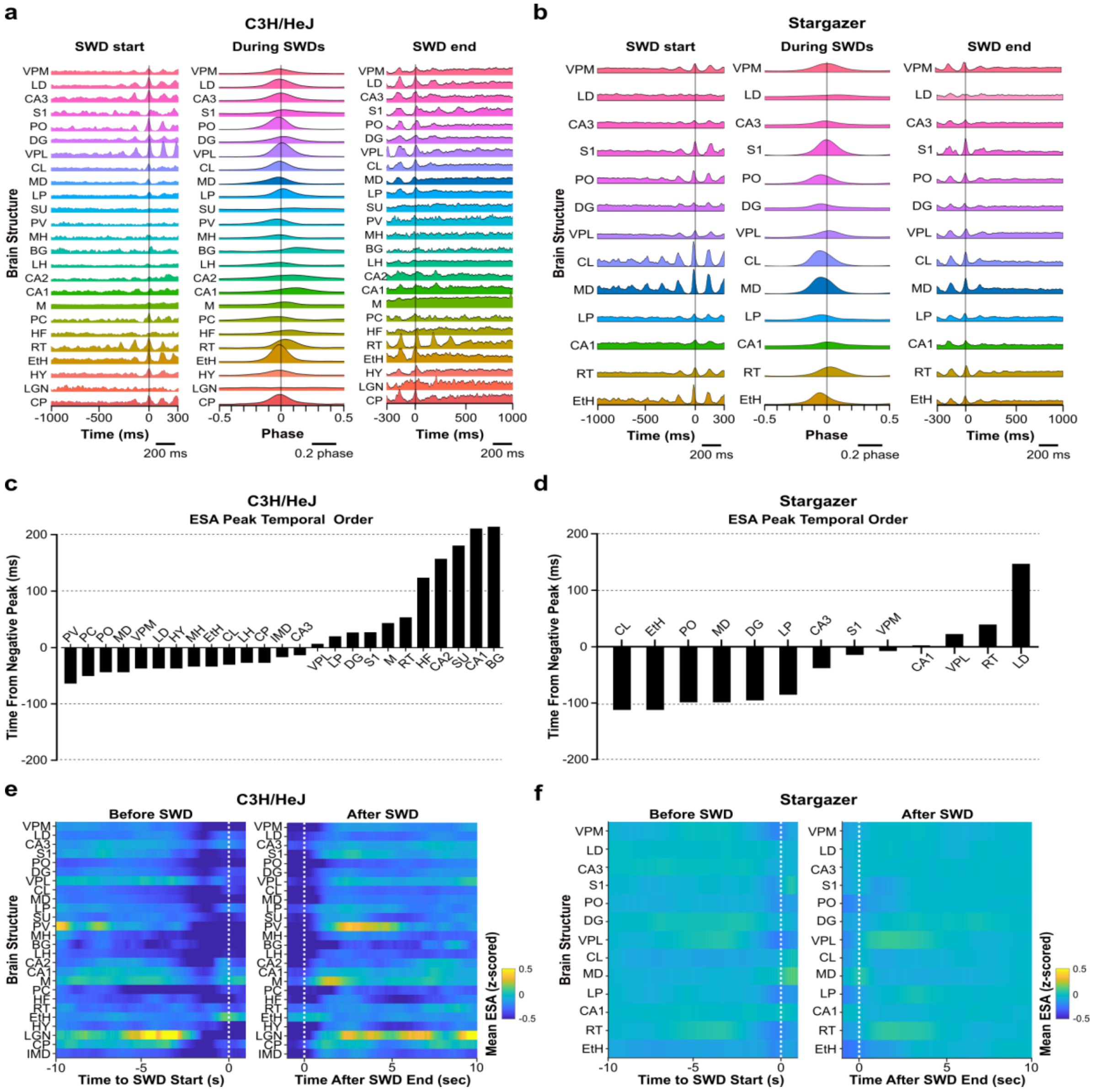


**Figure S7. ESA to resolve the temporal order of neuronal activation before, during, and after SWDs in C3H/HeJ and Stargazer mice**, Related to Figure 4. ***a***. Left: average ESA across multiple brain structures leading up to the SWD (0 seconds is SWD start) in C3H/HeJ mice. Middle: average ESA relative to negative peaks during SWDs. Right: average ESA relative to the end of SWDs (0 seconds is seizure end). ***b***. Comparable ESA before, during, and after SWDs for Stargazer mice. ***c***. Temporal order of ESA peaks relative to SWD negative peaks across brain structures. Negative values indicate ESA peaks occur before the cortical SWD negative peak, whereas positive values indicate ESA peaks occur after the negative SWD peak. ***d***. Temporal order of ESA peaks relative to SWD for Stargazer mice. ***e***. ESA leading up to the start of the C3H/HeJ SWD (left) and then after the SWD (right) over a longer timescale (-10 to +2 seconds from SWD start; -2 to +10 seconds from seizure end). We observed a marked decrease in ESA across most structures about 1 second before the SWD start. We also observed a noticeable return of ESA immediately after the SWD end. ESA traces in this panel were further temporally smoothed by calculating the moving mean over 1-second windows. ***f***. ESA leading to SWD start and after SWD end for Stargazer mice.

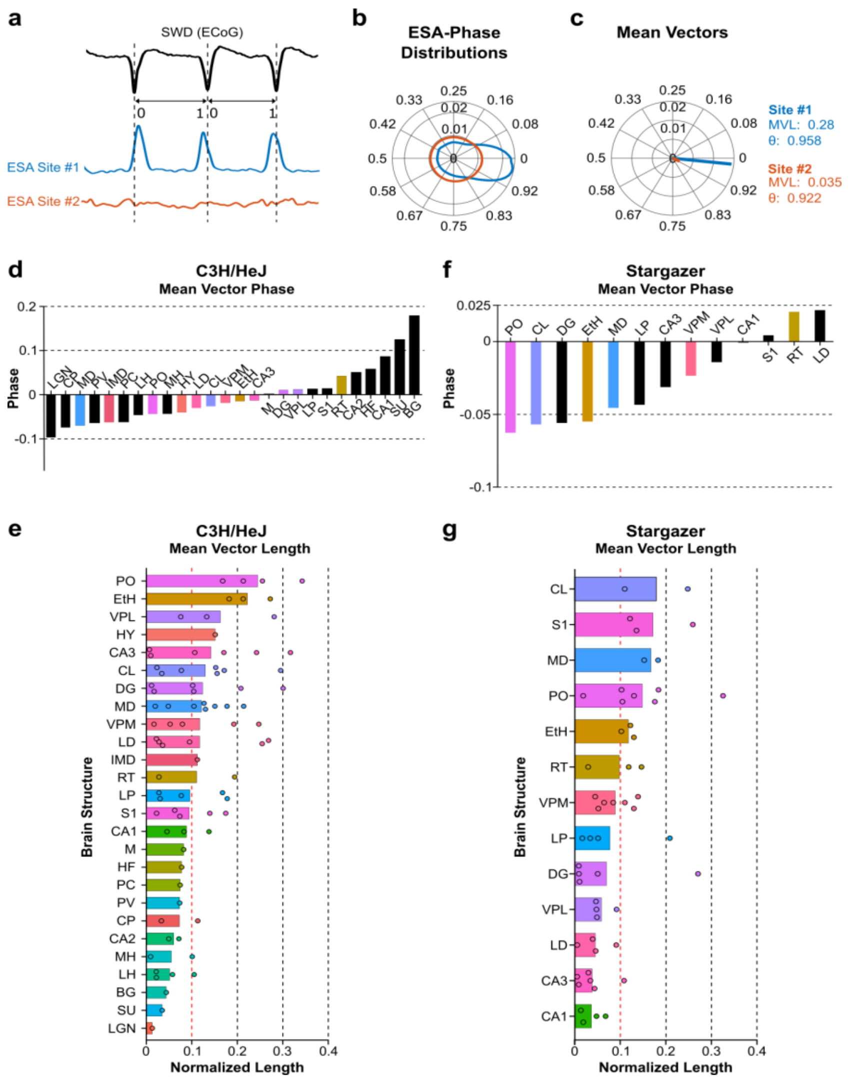


**Figure S8. Resolving SWD – ESA phase-amplitude coupling during SWDs using mean vectors**, Related to Figure 4. ***a***. Black, top: An example SWD recorded at the cortical surface. Each cycle of the SWD is divided into evenly spaced bins from 0 to 1. Blue, middle: ESA recorded at one electrode showing activity that is strongly phase-locked to the spike component of the SWD. Orange, bottom: ESA at a different electrode showing little to no phase-locking. ***b***. Normalized circular ESA phase distributions of the ESA in panel a. One example of ESA from a single electrode characterized by strong phase-locking is shown in blue. A second example showing weak phase-locking is presented in orange. ***c***. The corresponding mean vectors for normalized circular ESA phase distributions in b. Site #1 has a mean vector length of 0.28, indicating strong phase-locking, and an angle (θ) of 0.958. Site #2 has a mean vector length of 0.035, indicating little to no phase-locking, and an angle (θ) of 0.922. ***d***. Recorded brain regions sorted from left to right according to their ESA-SWD phase mean vector angles. Negative values indicate mean vector angles occur before the negative peaks in the SWD, as recorded at the cortical surface; positive values indicate angles that occur after the negative peaks. Colored bars indicate brain structures with mean vector lengths > 0.1, representing strong phase preference. Black bars indicate mean vector lengths below 0.1, representing weak phase preference (based on panel e). ***e***. Mean vector lengths in various recorded brain regions. Circles indicate the mean vector lengths from brain regions in individual mice. Bars represent the average across mice. ***f***. ESA-SWD phase mean vector angles for Stargazer mice. ***g***. Mean vector lengths for Stargazer mice.

**Table S1. Brain structures, abbreviations, and sample sizes**, Related to Figure 1. Brain structure names, abbreviations, and the number of single units and mice recorded from each structure. A dash (-) denotes structures not recorded in the Stargazer mouse.

| Brain Structure | Abbreviation | C3H/HeJ Mouse | | Stargazer Mouse | |
|---|---|---|---|---|---|
| | | # single units | # mice | # single units | # mice |
| Basal ganglia | BG | 4 | 1 | - | - |
| Caudate putamen | CP | 13 | 3 | - | - |
| Centrolateral thalamus | CL | 76 | 10 | 33 | 2 |
| Dentate gyrus | DG | 103 | 8 | 44 | 6 |
| Ethmoid nucleus of thalamus | EtH | 7 | 4 | 31 | 4 |
| Hippocampal formation | HF | 6 | 1 | - | - |
| Hippocampal CA1 | CA1 | 36 | 3 | 18 | 6 |
| Hippocampal CA2 | CA2 | 16 | 2 | - | - |
| Hippocampal CA3 | CA3 | 151 | 6 | 100 | 7 |
| Hypothalamus | HY | 6 | 1 | - | - |
| Intermediodorsal thalamus | IMD | 14 | 1 | - | - |
| Lateral dorsal thalamus | LD | 77 | 7 | 15 | 5 |
| Lateral habenula | LH | 50 | 9 | - | - |
| Lateral posterior nucleus of thalamus | LP | 54 | 8 | 66 | 6 |
| Medial habenula | MH | 3 | 3 | - | - |
| Mediodorsal thalamus | MD | 187 | 9 | 44 | 2 |
| Motor cortex | M | 53 | 1 | - | - |
| Paracentral nucleus | PC | 6 | 1 | - | - |
| Paraventricular thalamus | PV | 3 | 3 | - | - |
| Posterior complex of thalamus | PO | 58 | 6 | 88 | 8 |
| Primary somatosensory cortex | S1 | 25 | 5 | 135 | 4 |
| Reticular thalamus | RT | 10 | 6 | 64 | 3 |
| Subthalamic nucleus | SU | 6 | 1 | - | - |
| Ventral posteriolateral nucleus of thalamus | VPL | 17 | 5 | 102 | 6 |
| Ventral posteriomedial nucleus of thalamus | VPM | 61 | 6 | 106 | 7 |
| Lateral geniculate nucleus of thalamus | LGN | 1 | 1 | - | - |

**Table S2. Mean firing rates during non-seizures and SWDs**, Related to Figure 2. A dash (-) denotes structures not recorded in the Stargazer mouse and therefore mean firing rate (FR) was not calculated.

| Brain Structure | Abbreviation | C3H/HeJ Mouse Mean Firing Rate (Hz) | | Stargazer Mouse Mean Firing Rate (Hz) | |
|---|---|---|---|---|---|
| | | Non-seizures | SWDs | Non-seizures | SWDs |
| Basal ganglia | BG | 45.19 | 53.05 | - | - |
| Caudate putamen | CP | 27.61 | 22.96 | - | - |
| Centrolateral thalamus | CL | 7.84 | 6.13 | 4.42 | 4.58 |
| Dentate gyrus | DG | 7.18 | 5.65 | 3.77 | 2.88 |
| Ethmoid nucleus of thalamus | EtH | 4.20 | 5.06 | 4.27 | 3.90 |
| Hippocampal formation | HF | 4.28 | 3.51 | - | - |
| Hippocampal CA1 | CA1 | 7.69 | 6.56 | 13.14 | 13.08 |
| Hippocampal CA2 | CA2 | 5.59 | 5.68 | - | - |
| Hippocampal CA3 | CA3 | 7.96 | 5.67 | - | - |
| Hypothalamus | HY | 8.20 | 6.83 | 8.76 | 8.21 |
| Intermediodorsal thalamus | IMD | 2.80 | 2.18 | - | - |
| Lateral dorsal thalamus | LD | 7.80 | 4.64 | 16.25 | 15.71 |
| Lateral habenula | LH | 10.61 | 7.54 | - | - |
| Lateral posterior nucleus of thalamus | LP | 5.57 | 5.95 | 3.94 | 2.43 |
| Medial habenula | MH | 12.29 | 11.89 | - | - |
| Mediodorsal thalamus | MD | 10.05 | 7.39 | 3.95 | 4.35 |
| Motor cortex | M | 4.77 | 3.00 | - | - |
| Paracentral nucleus | PC | 10.78 | 7.25 | - | - |
| Paraventricular thalamus | PV | 6.47 | 6.42 | - | - |
| Posterior complex of thalamus | PO | 8.12 | 6.49 | 5.56 | 3.94 |
| Primary somatosensory cortex | S1 | 3.44 | 1.94 | 4.37 | 3.99 |
| Reticular thalamus | RT | 11.34 | 4.36 | 11.23 | 10.63 |
| Subthalamic nucleus | SU | 8.00 | 6.51 | - | - |
| Ventral posteriolateral nucleus of thalamus | VPL | 15.20 | 15.28 | 7.10 | 5.24 |
| Ventral posteriomedial nucleus of thalamus | VPM | 11.61 | 7.50 | 6.23 | 4.99 |
| Lateral geniculate nucleus of thalamus | LGN | 6.15 | 4.23 | - | - |

**Table S3. Putative excitatory and inhibitory unit classification and SWD-related firing changes**, Related to Figure 2**.** Number and proportion of putative excitatory and inhibitory units are summarized for each recorded brain structure in C3H/HeJ and Stargazer mice. Units were classified based on extracellular waveform features. For each cell type, the table reports the fraction, and percentage of units exhibiting consistently increased firing during SWDs relative to baseline (mean z-score > 2 for at least 50% SWDs), referred to as responsive units ("Resp"). A dash (-) denotes lack of putative excitatory/inhibitory units recorded.

| | **C3H/HeJ mouse** | | | | **Stargazer mouse** | | | |
|---|---|---|---|---|---|---|---|---|
| | **# putative *excitatory* units** | | **# putative *inhibitory* units** | | **# putative *excitatory* units** | | **# putative *inhibitory* units** | |
| **Abbr.** | **Resp / total *exc* units** | **% Resp *exc* units** | **Resp / total *inh* units** | **% Resp *inh* units** | **Resp / total *exc* units** | **% Resp *exc* units** | **Resp / total *inh* units** | **% Resp *inh* units** |
| BG | 0 / 2 | 0% | 2 / 2 | 100% | - | | | |
| CP | 0 / 5 | 0% | 0 / 2 | 0% | - | | | |
| CL | 13 / 63 | 21% | - / 0 | | 0 / 32 | 0% | - / 0 | - |
| DG | 10 / 80 | 13% | 1 / 5 | 20% | 0 / 32 | 0% | 0 / 2 | 0% |
| EtH | 1 / 5 | 20% | 0 / 1 | 0% | 0 / 23 | 0% | 0 / 4 | 0% |
| HF | 0 / 2 | 0% | - / 0 | | - | | - / - | - |
| CA1 | 2 / 27 | 7% | 0 / 2 | 0% | 0 / 7 | 0% | 0 / 6 | 0% |
| CA2 | 6 / 14 | 43% | - / 0 | | - | | - / - | - |
| CA3 | 15 / 97 | 15% | 1 / 12 | 8% | 0 / 67 | 0% | 0 / 12 | 0% |
| HY | 0 / 4 | 0% | - / 0 | | - | | - / - | - |
| IMD | 0 / 14 | 0% | - / 0 | | - | | - / - | - |
| LD | 8 / 54 | 15% | 1 / 3 | 33% | 0 / 11 | 0% | 0 / 1 | 0% |
| LH | 4 / 49 | 8% | 0 / 1 | 0% | - | | - / - | - |
| LP | 11 / 51 | 22% | - / 0 | | 0 / 53 | 0% | 0 / 1 | 0% |
| MH | 0 / 3 | 0% | - / 0 | | - | | - / - | - |
| MD | 8 / 147 | 5% | - / 0 | | 0 / 43 | 0% | - / 0 | - |
| PC | - / - | | - / - | | - | | - / - | - |
| M | 1 / 42 | 2% | 0 / 8 | 0% | - | | - / - | - |
| PV | 0 / 3 | 0% | - / 0 | | - | | - / - | - |
| PO | 7 / 57 | 12% | - / 0 | | 0 / 63 | 0% | 0 / 3 | 0% |
| S1 | 1 / 18 | 6% | 1 / 3 | 33% | 4 / 96 | 4% | 0 / 7 | 0% |
| RT | 0 / 3 | 0% | 0 / 4 | 0% | 0 / 3 | 0% | 0 / 53 | 0% |
| SU | 0 / 2 | 0% | 0 / 2 | 0% | - / - | | - / - | - |
| VPL | 0 / 6 | 0% | 2 / 5 | 40% | 0 / 36 | 0% | 6 / 31 | 19% |
| VPM | 1 / 44 | 2% | 0 / 5 | 0% | 0 / 53 | 0% | 3 / 22 | 14% |
| LGN | - / 0 | - | - / 0 | | - / - | | - / - | - |

**Table S4. Electrode tip coordinates for midline thalamic stimulation**, Related to Figure 6. Stereotaxic coordinates of left and right electrode tips; anteroposterior (AP), mediolateral (ML), and dorsoventral (DV), in mm, for each animal, determined by post hoc histological verification. Animals with off-target electrode placements (underlined coordinates) were excluded from analysis and are indicated accordingly in the final column.

| **Mouse** | **Hemisphere** | **Electrode tip coordinate** | | | **Inclusion** |
|---|---|---|---|---|---|
| | | AP | ML | DV | |
| **C3H/HeJ #1** | *Left* | -1.4 | 0.5 | 2.8 | Included |
| | *Right* | -1.4 | -0.6 | 2.8 | |
| **C3H/HeJ #2** | *Left* | -2.1 | 0.8 | 2.4 | Excluded |
| | *Right* | -2.6 | -0.8 | 2.8 | |
| **C3H/HeJ #3** | *Left* | -2.1 | 0.7 | 3.0 | Included |
| | *Right* | -2.1 | -0.8 | 3.0 | |
| **C3H/HeJ #4** | *Left* | -1.7 | 0.9 | 3.2 | Included |
| | *Right* | -1.5 | -0.7 | 3.2 | |
| **C3H/HeJ #5** | *Left* | -1.3 | 0.8 | 3.5 | Included |
| | *Right* | -1.3 | -0.8 | 3.5 | |
| **C3H/HeJ #6** | *Left* | -1.5 | 0.3 | 3.4 | Included |
| | *Right* | -1.3 | -0.4 | 2.9 | |
| **Stargazer #1** | *Left* | -1.8 | 0.5 | 2.6 | Included |
| | *Right* | -2.1 | -0.4 | 2.0 | |
| **Stargazer #2** | *Left* | -1.7 | 0.6 | 2.8 | Included |
| | *Right* | -1.6 | -0.4 | 2.6 | |
| **Stargazer #3** | *Left* | -2.0 | 0.5 | 2.7 | Included |
| | *Right* | -1.8 | -0.8 | 3.2 | |
| **Stargazer #4** | *Left* | -1.7 | 0.4 | 3.0 | Excluded |
| | *Right* | -2.6 | -0.3 | 2.3 | |
